# Highly similar sequence and structure yet different biophysical behaviour: A computational study of two triosephosphate isomerases

**DOI:** 10.1101/2021.10.13.464197

**Authors:** Cecilia Chávez-García, Mikko Karttunen

## Abstract

Homodimeric triosephosphate isomerases (TIM) from Trypanosoma cruzi (TcTIM) and Trypanosoma brucei (TbTIM) have a markedly similar amino acid sequences and three-dimensional structures. However, several of their biophysical parameters, such as their susceptibility to sulfhydryl agents and their reactivation speed after being denatured, have significant differences. The causes of these differences were explored with microsecond-scale molecular dynamics (MD) simulations of three different TIM proteins: TcTIM, TbTIM and a chimeric protein, Mut1. We examined their electrostatic interactions and explored the impact of simulation length on them. The same salt bridge between catalytic residues Lys 14 and Glu 98 was observed in all three proteins but key differences were found in other interactions that the catalytic amino acids form. In particular, a cation-π interaction between catalytic amino acids Lys 14 and His 96, and both a salt bridge and a hydrogen bond between catalytic Glu168 and residue Arg100, were only observed in TcTIM. Furthermore, although TcTIM forms less hydrogen bonds than TbTIM and Mut1, its hydrogen bond network spans almost the entire protein, connecting the residues in both monomers. This work provides new insight on the mechanisms that give rise to the different behaviour of these proteins. The results also show the importance of long simulations.

## INTRODUCTION

One of the most common structural motifs in proteins is the triosephosphate isomerase (TIM) barrel, which is present in ~10% of all known proteins and is the most common enzyme fold in the Protein Data Bank (PDB) database^1–4^. TIM is an enzyme which takes part in the fifth step of glycolysis by interconverting glyceraldehyde 3-phosphate into dihydroxyacetone phosphate. The TIM barrel consists of an eightfold repeat of (βα) units in such a way that β-strands in the inside are surrounded by α-helices on the outside. TIM is present in almost all organisms and is usually found as a dimer, although it can form a tetramer in some extremophile organisms^5–8^. It is completely active only in the dimeric form even though each monomer contains the catalytic residues (N12, K14, H96 and E168). The catalytic residues are strictly conserved throughout the whole TIM family^1,9,10^. TIM is essential for maintaining life under anaerobic conditions and, consequently, it has been used as a target for drug design when dealing with human parasites^11–13^.

There are multiple instances in which homologous enzymes with high similarity have significant differences in their biophysical parameters^14–17^. This is exemplified by the triosephosphate isomerases of *Trypanosoma cruzi* (TcTIM), the parasite that causes Chagas’ disease, and *Trypanosoma brucei* (TbTIM), causative agent of the African sleeping sickness. They have a sequence identity of 73.9% and a sequence similarity of 92.4%^18^ (Fig. 1). Previous works have found significant differences in their susceptibility to sulfhydryl agents, reactivation speed after being denatured with chemical agents such as guanidine hydrochloride, and their proteolysis susceptibility with subtilisin^16,18–20^. Interestingly, a study of TcTIM and TbTIM by Rodríguez-Bolaños *et al.*^21^ showed that it is sufficient to mutate 13 amino acids on TbTIM to obtain TcTIM-like behaviour in reactivation experiments. Circular dichroism indicated that the chimeric proteins had the same fold as the native, however, the role that these mutations have on the structure and dynamics of the proteins is not well understood.

**Figure 1.**
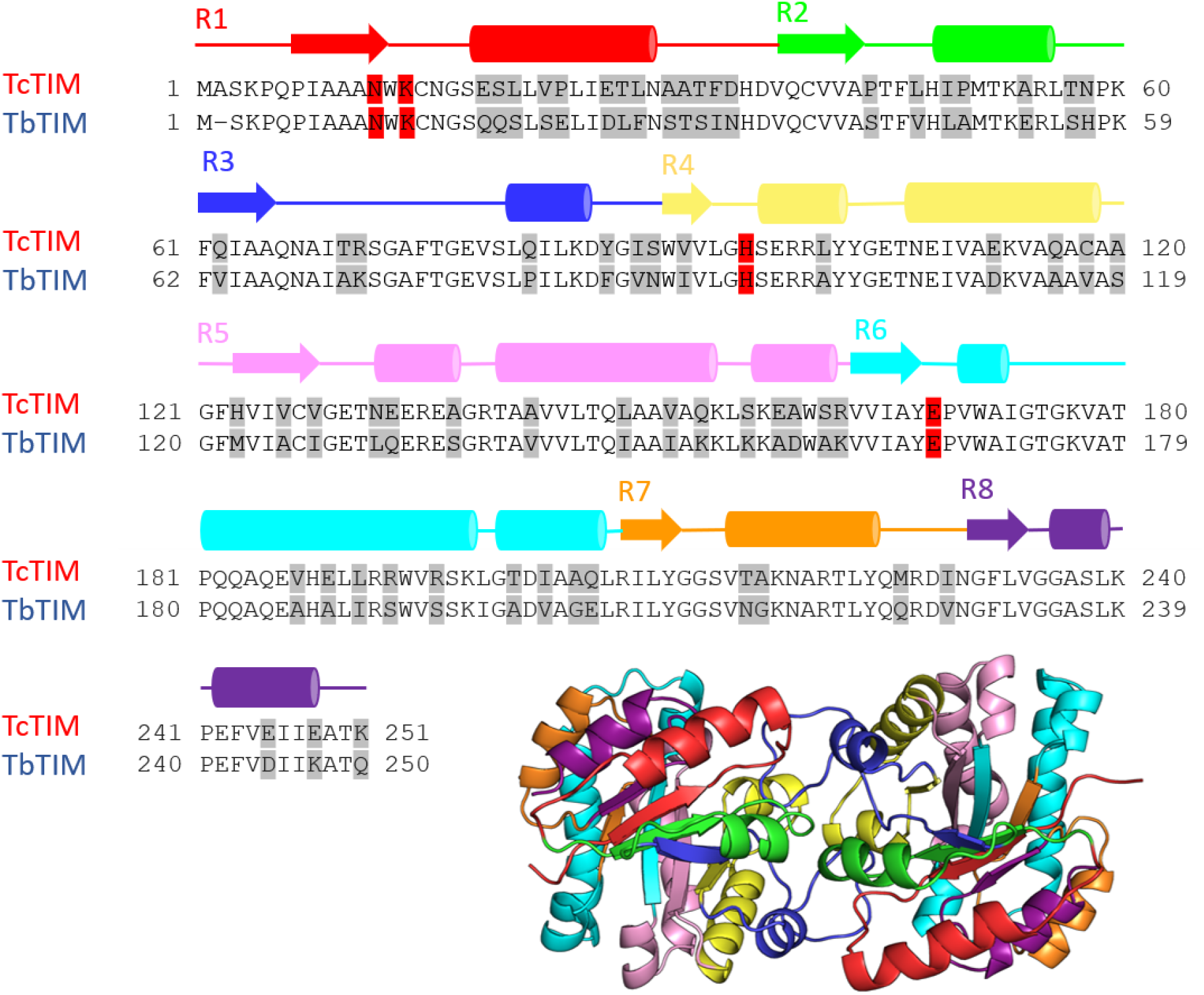
Alignment of TcTIM and TbTIM sequences with the secondary structure elements marked as lines (loops), β-strands (arrows) and α-helices (cylinders). The colors in each motif correspond to the 3D structure below it. The differences in the amino acids are highlighted in gray and the catalytic residues in red.

For many years, studies have focused on the interaction between TIM proteins and benzothiazoles, which have been found to deactivate the enzyme^13,22,23^. While there is a plethora of experimental data, there are very few MD simulations of TcTIM and TbTIM in the absence of ligands. Some of the simulations are only 40-60 ns long, and thus cannot capture phenomena that occur in longer timescales.^24–26^ There is only one study in the microsecond scale. In that Dantu and Groenhof use a combination of QM/MM and crystal unit cell simulations^27^. However, this study focuses on the effect of binding of substrates in loops 5, 6 and 7. The most comprehensive study on TIM so far used conventional MD and enhanced sampling techniques to characterize the motion of loops 6 and 7^28^. This study showed that loop 6 does not follow a simple two-state rigid-body transition as previously thought. However, it did not explore in detail the interactions between the two monomers. The simulations that report the root-mean-square fluctuation (RMSF) of TIM proteins have consistently found the highest RMSF values in loops 5 and 6^22,24,25^, which is in good agreement with our work.

Here we present microsecond-scale all-atom simulations of TcTIM, TbTIM and a chimeric TbTIM with the 13 mutations as identified by Rodríguez-Bolaños *et al*^21^. A conserved salt bridge between catalytic residues Lys 14 and Glu 98^29,30^ was observed in all three proteins. In contrast, a cation-π interaction between catalytic amino acids Lys 14 and His 96, and both a salt bridge and a hydrogen bond between catalytic Glu 168 and Arg 100 were only observed in TcTIM. Furthermore, TcTIM and TbTIM exhibited different hydrogen bond networks, with the chimeric protein behaving similar to TbTIM. The hydrogen bond network observed in these proteins helps to explain why regions 1, 4 and 8 become more rigid when the dimer is formed.

## METHODS

All-atom MD simulations were performed on three TIM proteins: 1) the TIM from *T. brucei* (TbTIM), 2) the TIM from *T. cruzi* (TcTIM), and 3) a chimeric protein: [TcTIM:2; TbTIM:1,3-8; Q18E, E23P, D26E, S32T, I33F, N34D] henceforth referred to as Mut1. Each protein was simulated as a dimer. The initial structures were taken from the Protein Data Bank (PDB ids: 1TCD^31^ and 5TIM^32^) and any missing side chains were completed using the whatif web server^33^. Mut1 was built using TbTIM as template and mutating the required amino acids using Pymol^34^.

Each protein was placed in a dodecahedral box in which the distance from the edges of the box to every atom in the protein was at least 1 nm. The box was solvated with explicit water and 150 mM of NaCl was added to reproduce physiological conditions. Counterions were added to keep the overall charge neutrality of the systems, 6 Cl^−^ ions for the TcTIM and 10 Cl^−^ ions for both 5TIM and Mut1. All simulations were performed using GROMACS 2016.3^35^ with the TIP3P water model^36^ and the CHARMM36m force field^37^.

Each system was first energy minimized using the method of steepest descents and pre-equilibrated at constant particle number, temperature and volume, for 100 ps. The pre-equilibration was followed by a production run with a time step of 2 fs. The Lennard-Jones potential was truncated using a shift function between 1.0 and 1.2 nm. Electrostatic interactions were calculated using the particle-mesh Ewald method (PME)^38,39^ with a real space cut-off of 1.2 nm. The temperature was set to 310 K with the V-rescale algorithm^40^ and pressure was kept at 1 atm using the Parrinello-Rahman barostat^41^. Bonds involving hydrogens were constrained using the Parallel Linear Constraint Solver (P-LINCS) algorithm^42^. The root-mean-square deviation (RMSD) was used to monitor equilibration. Since the RMSD for TcTIM kept increasing during the first microsecond, the simulations were extended to 3 μs. We will return to this issue later in Results.

Trajectory analyses were performed using Gromacs built-in tools^35^, MDAnalysis^43,44^ and the VMD plug-ins Salt bridges^45^ and RIP-MD^46^. RIP-MD generates residue interaction networks (RINs) from MD trajectory files. In a RIN, the nodes of the network represent amino-acid residues and the connections between them depict non-covalent interactions. These include hydrogen bonds, salt bridges, cation-π, π–π, arginine–arginine, and Coulomb interactions. RIP-MD starts with a MD trajectory and the parameters defining the interactions as input. It then searches for interactions between all atoms in each snapshot of the trajectory. Finally, it generates a consensus RIN where edges exist if they are present in at least a given percentage of the snapshots. For our study, we used a 30% threshold.

## RESULTS

We performed MD simulations on three different TIM proteins: TcTIM, TbTIM and Mut1. Figure 2 shows that the RMSD of all three systems increase throughout the first 1500 ns, after which they stabilized. For this reason, the simulations were extended to 3 μs and all reported averages were calculated during the last microsecond of the trajectories.

**Figure 2.**
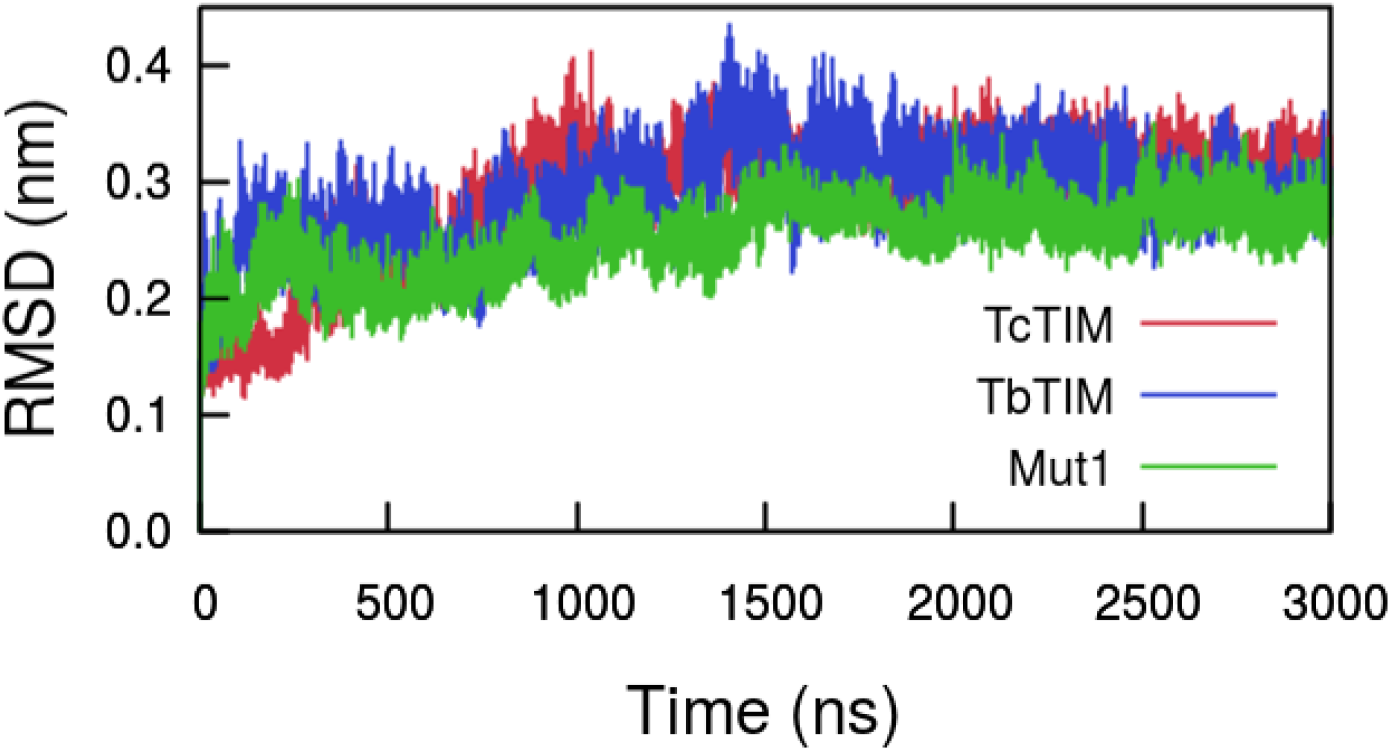
Root-mean-square deviation with respect to the crystal structure. RMSD increases throughout the first 1500 ns before stabilizing.

The root-mean-square fluctuations (RMSF) of most of the residues in the three proteins are very similar (Fig. 3). The main differences appear at the highest peaks, which are located at residues 133-137 (loop 5) and 173-178 (loop 6) in both monomers. The amino acids in loop 6 correspond to the catalytic loop. This loop has a “phosphate gripper” motif^47^ which is likely engaged in substrate binding and product release, as its opening and closing motion has a rate constant that closely matches the turnover time for catalysis^48,49^. TcTIM is the only protein with peaks at loops 5 and 6 in monomer A and TbTIM is the only protein with a peak in loop 5 monomer B. All three proteins have a peak in loop 6 monomer B, albeit the peak in TbTIM is almost two times larger than in the other two systems. TcTIM is the only protein in which the first residues of monomer B have a very low RMSF value, indicating an interaction with monomer A.

**Figure 3.**
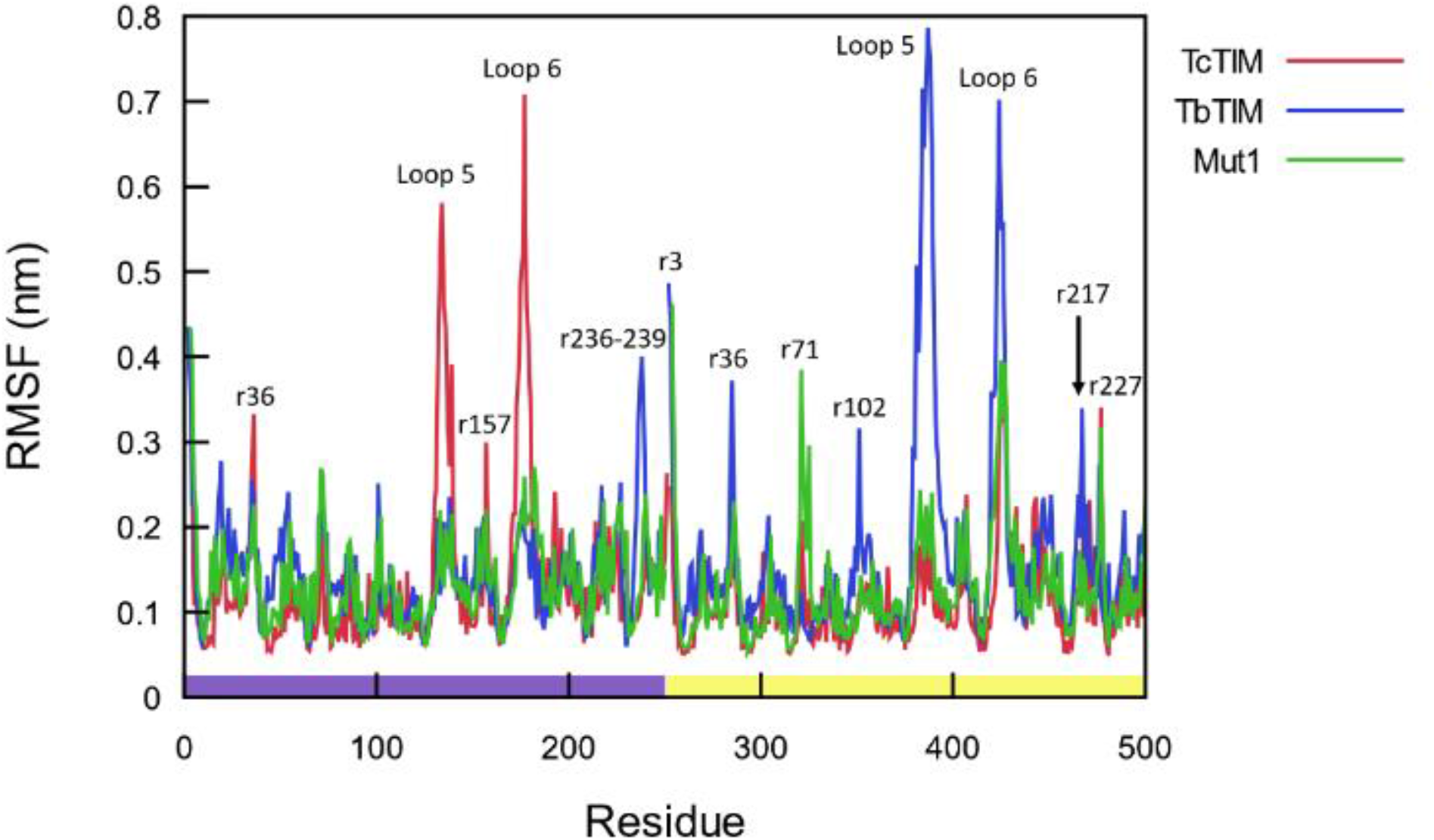
Root mean square fluctuations for the last microsecond. Since each protein was simulated as a dimer, the color bar at the bottom distinguishes the residues in monomer A (purple) from those in monomer B (yellow). The main peaks are located at loops 5 and 6 in each monomer.

In a previous computational study of TcTIM^25^, loops 5 and 6 were reported to have the largest fluctuations. This study used GROMOS96 (43a2)^50^, a united-atom force field and the SPC water model^51^. The fact that the same results were obtained with two very different force fields (GROMOS and CHARMM) underlines their robustness and their independence of the chosen force field. All the other main peaks correspond to amino acids located in loops with the exception of residues 236-239, which span the short helix in region 8.

The number of contacts between monomers, defined as amino acids whose Cβ atoms (Cα for glycine) are within 0.8 nm distance, is shown in Figure 4. Even though a protein residue-residue contact is not uniquely defined, this definition captures all possible interactions between two residues and it has been used in a number of previous studies^52–55^. The number of contacts changes for both TcTIM and Mut1 during the first microsecond before stabilizing. In contrast, the number of contacts between TbTIM’s monomers fluctuated throughout the entire simulation. The average number of contacts between monomers during the last microsecond is 119 for TcTIM, 89 for TbTIM and 116 for Mut1. This is in good agreement with previous experiments, since the number of contacts between monomers can be related to its thermal stability and TcTIM has higher thermal stability than TbTIM^21^.

**Figure 4.**
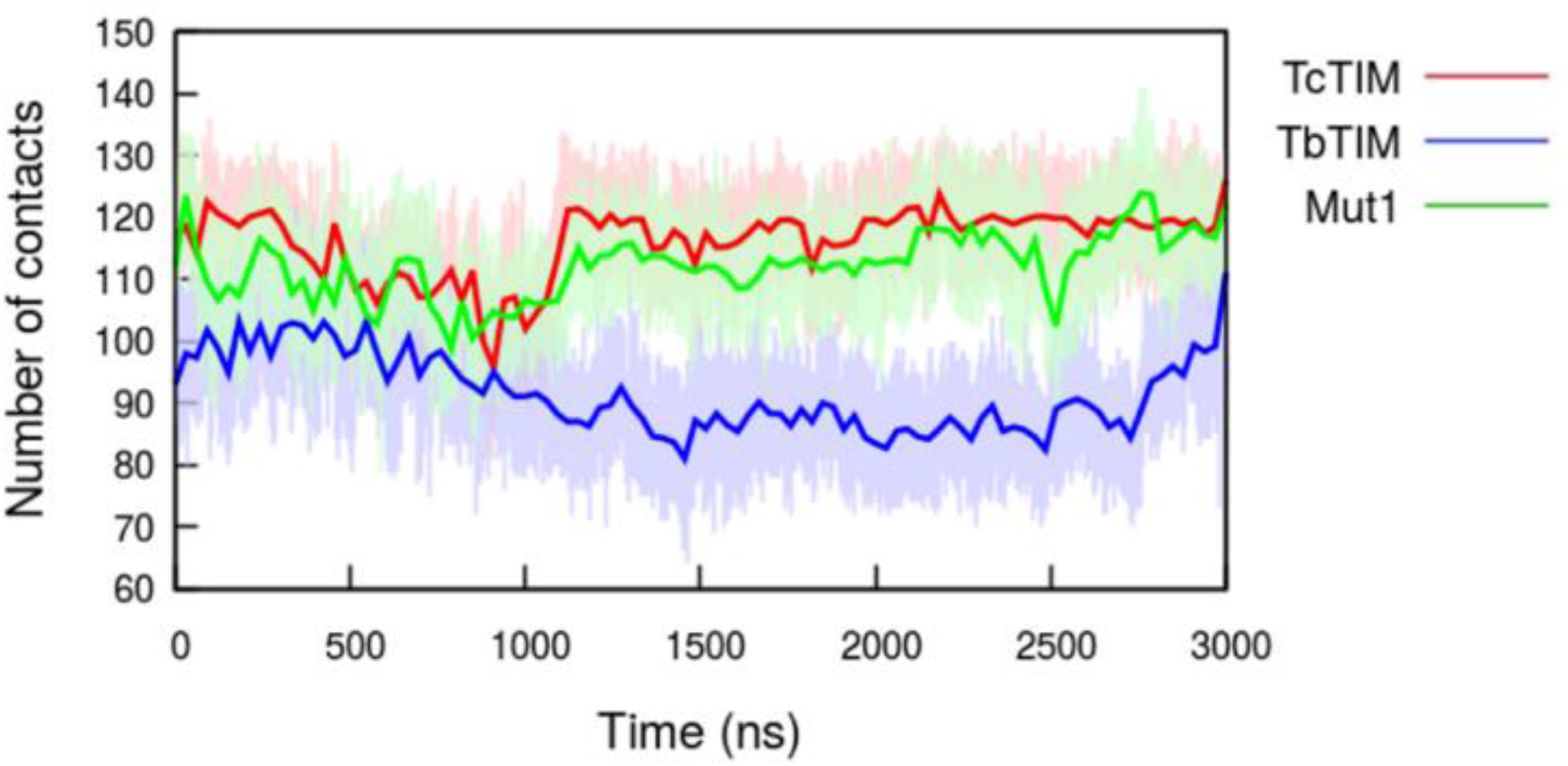
Number of contacts between monomers. Residues are considered to be in contact if their respective Cβ atoms (Cα for glycine) are less than 0.8 nm apart. The number of contacts in TcTIM and Mut1 decreases over the first microsecond, increases over the next 200 ns and then stabilizes. In TbTIM, they decrease during the first half of the simulation and increase again after 1500 ns. Solid lines are Bézier curves that interpolate the data.

In order to identify if the systems were in the open or closed conformations, the minimum distance between loop 6 (residues 170-180) and loop 7 (residues 211-216) was measured (Fig. 5, S1). Five different states were sampled in TcTIM, with distances 0.18, 0.28, 0.50, 0.67 and 0.80 nm. Only the first three states were observed in TbTIM and Mut1. It was previously reported that loop 6 can sample multiple conformational states with the tip of the loop moving ~0.7 nm between the fully open and fully closed conformations^28^. This is in good agreement with the difference between the two extremes observed in the current TcTIM simulation. TbTIM fluctuated the most between states, and while Mut1 also showed many fluctuations, its loop in monomer B remained in the fully closed conformation for the last microsecond of the simulation. The cross-correlation between the open-close conformation of the two monomers decayed to zero during the simulation, indicating that the movement of these loops is independent between monomers.

**Figure 5.**
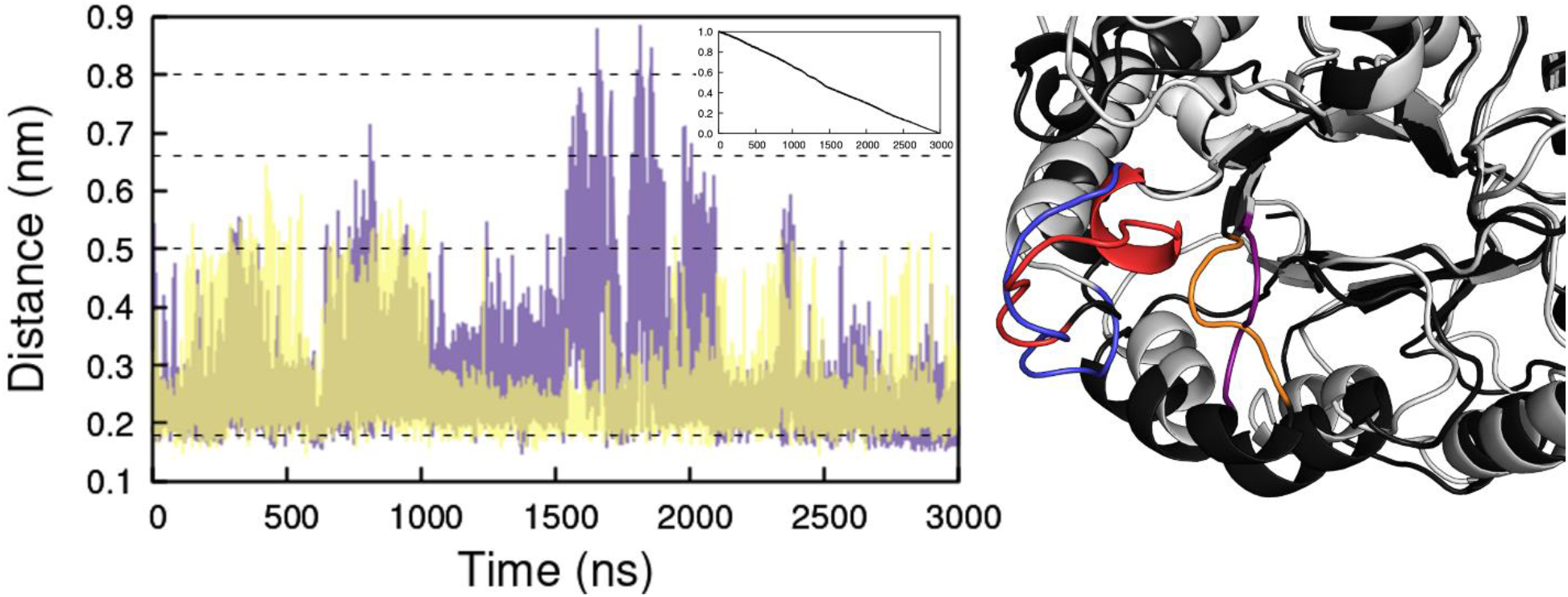
Minimum distance between loops 6 and 7 in TcTIM for monomer A (purple) and monomer B (yellow). Dashed lines mark the five different states sampled by the loops. Inset: cross-correlation between the loop state of the two monomers. Right: alignment of the open (gray) and closed (black) conformations. Loop 6 is shown in red for the closed conformation and in blue for the open conformation, loop 7 is shown in orange (closed) and purple (open).

### Electrostatic interactions

We used the RIP-MD^46^ plugin for VMD^45^ to analyze the last 2 μs of each trajectory and to compute the following electrostatic interactions between all residues: salt bridges, cation-π interactions, π-π interactions and hydrogen bonds. No arginine-arginine interactions were found in the simulations. We will describe these interactions in the section below in more detail, but Table 1 summarizes the parameters defining them.

**Table 1.** Summary of interactions defined in RIP-MD.

|  |  |  |
| --- | --- | --- |
| **Hydrogen bonds** | $\text{dist}(\text{donor}, \text{acceptor}) \leq d$ | $d = 3 \text{ \AA}$ |
| | $\theta(\overrightarrow{C-H}, \overrightarrow{\text{acceptor}}) \geq a$ | $a = 120^\circ$ |
| **Salt bridges** | Contacts between NH/NZ groups of ARG/LYS<br>and OE/OD in ASP/GLU $\leq d$ | $d = 6 \text{ \AA}$ |
| **Cation-π** | $\text{dist}(\text{aromatic ring}, \text{cation}) \leq d$ | $d = 6 \text{ \AA}$ |
| **interactions** | $\theta(\overrightarrow{\text{normal vector ring}}, \overrightarrow{\text{ring center} - \text{cation}}) = a$ | $a \in [0^\circ, 60^\circ]$ or |
| | | $a \in [120^\circ, 180^\circ]$ |
| **π-π interactions** | $\text{dist}(\text{aromatic ring}, \text{aromatic ring}) \leq d$ | $d = 7 \text{ \AA}$ |
| **Arg-Arg** | $\text{dist}(\text{guanidine}, \text{guanidine}) \leq d$ | $d = 5 \text{ \AA}$ |

From the three proteins, only TcTIM had cation-π interactions (Fig. S2). These interactions were defined between the geometric center of the ring in the aromatic residue and the charged atom in the second residue, with a cutoff distance of 0.6 nm. This threshold was chosen because 99% of significant cation-pi interactions occur within a distance of 0.6 nm^56^. The only cation-π interaction between monomers occurs between amino acids Tyr 103 in monomer A and Arg 99 in monomer B. Figure 6 shows the distance that defines this cation-π interaction throughout the simulation. The distance fluctuates throughout time but remains within the limits that define the cation-π interaction during most frames in the last 1,500 ns of the simulation.

**Figure 6.**
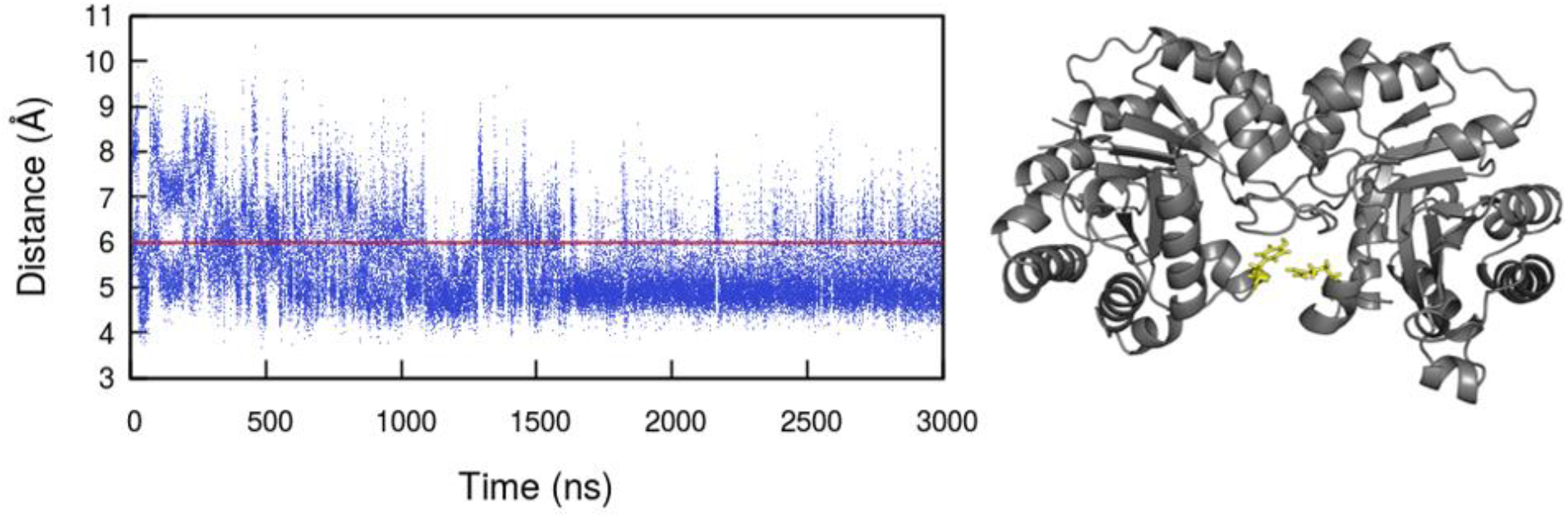
Cation- π interaction between Tyr 103 in monomer A and Arg 99 in monomer B in TcTIM. This interaction is defined by the distance between the geometric center of the aromatic residue in tyrosine and the charged atom in arginine, with a cutoff distance of 0.6 nm. This interaction was only observed in TcTIM. Right: these amino acids are located in region 4. A red line marks the threshold that defines the cation-π interaction.

Π-π interactions were defined with the distance between the geometric centers of the rings in the aromatic amino acids, with a cutoff distance of 0.7 nm. The distance between two interacting aromatic rings is geometry dependent and varies between 0.45 and 0.7 nm^57^. All three systems presented π-π interactions (Fig. S3). Out of the 14 π-π interactions in TcTIM, four occurred at the interface between monomers. TbTIM only presented five π-π interactions and the mutant had four, two of which occurred at the interface. Interestingly, two of the π-π interactions in the mutant correspond to interactions in TcTIM and the other two to interactions in TbTIM.

### Residue interaction networks

In RIP-MD^46^, salt bridges are treated as a contact between two heavy atoms of opposite charge with a distance threshold of 0.6 nm^46,58^. The same salt bridges were observed in the three proteins: between Glu 78 (Glu 77 in TbTIM) and Arg 99 (Arg 98 in TbTIM) of both monomers (Fig. S4). Residue Arg 55 forms a salt bridge with residue 27 from the same monomer, in both monomers in TcTIM. Since RIP-MD^46^ does not provide the time dependence of the salt bridges, we used the Salt Bridge VMD plugin^45^ to calculate them. This plugin uses a different cutoff distance (0.32 nm) to define a salt bridge, however, as long as the interacting atoms are within the threshold in one frame, the program outputs the distance between them as a function of time. Figure 7 illustrates the fluctuations of the distance between atoms that form salt bridges. The interaction between Glu 27 and Arg 55 in TcTIM fluctuates considerably and the salt bridge is defined in only a portion of the frames. In contrast, the salt bridge between Glu 98 and Lys 14 stays well within the limit that defines this interaction throughout the whole simulation.

**Figure 7.**
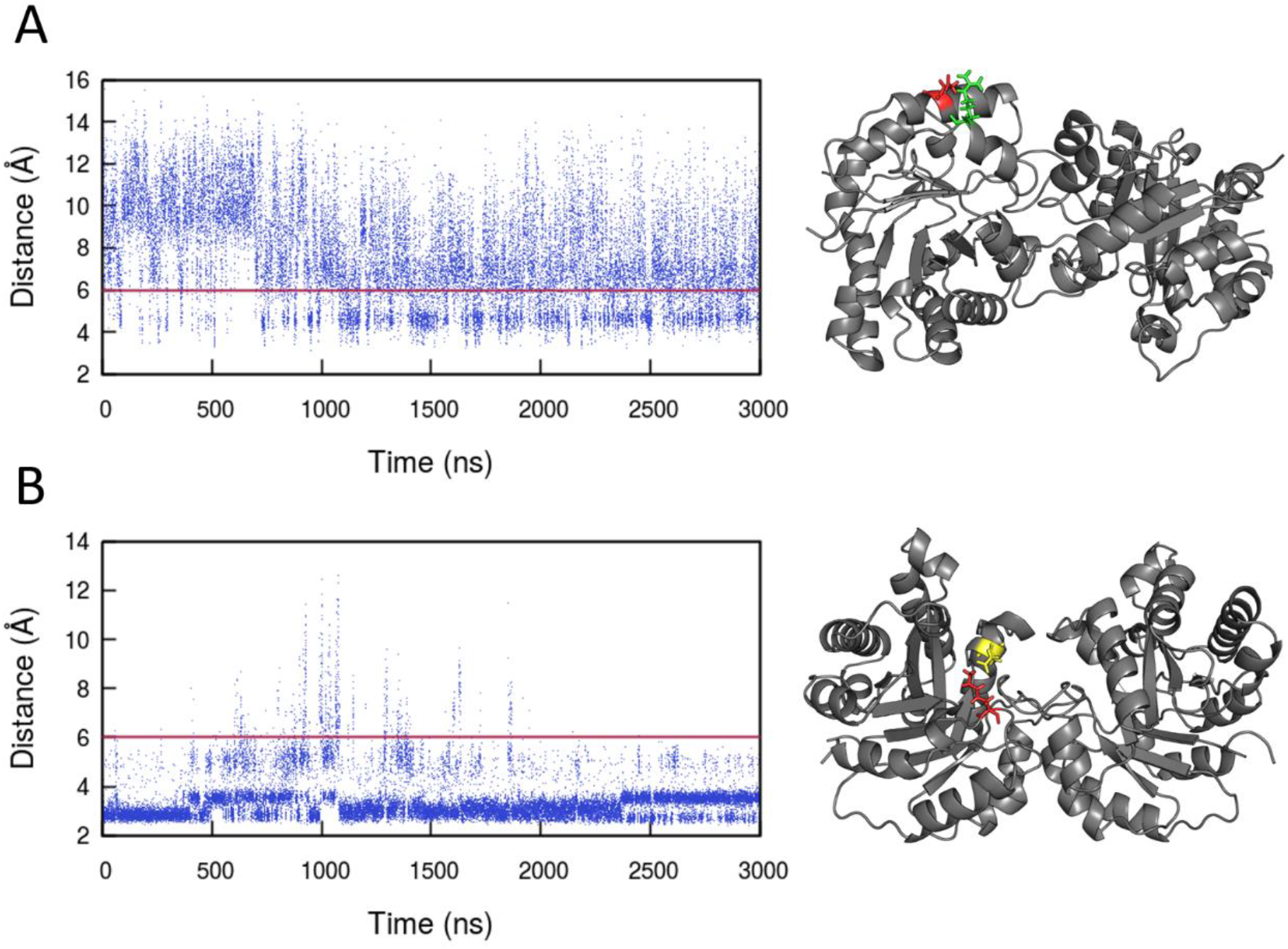
Salt bridges in TcTIM between Glu27 monomer A and Arg55 monomer A (A), and between Glu98 monomer A and Lys14 monomer A (B). These interactions were computed using the Salt Bridges plugin for VMD^45^. Right: 3D location of the amino acids. Residues Glu27 and Lys14 are shown in red (region 1), Arg55 in green (region 2) and Glu98 in yellow (region 4). A red line marks the threshold that defines the salt bridge interaction.

Hydrogen bonds were defined using a cutoff radius of 0.3 nm for atoms whose acceptor-hydrogen-donor angle is greater than 120°^59^. The average number of hydrogen bonds over the last microsecond was 345 for TcTIM and 369 ± 1% for TbTIM and Mut1 (Fig. 8). Figures S5-S15 show the hydrogen bond networks in the three proteins. For clarity, hydrogen bonds formed between neighbouring amino acids (less than 4 residues apart) have been removed from the graphs. Even though TcTIM forms less hydrogen bonds than TbTIM, they connect amino acids in a network that involve more interactions between monomers and extends throughout the whole protein (Fig. S16).

**Figure 8.**
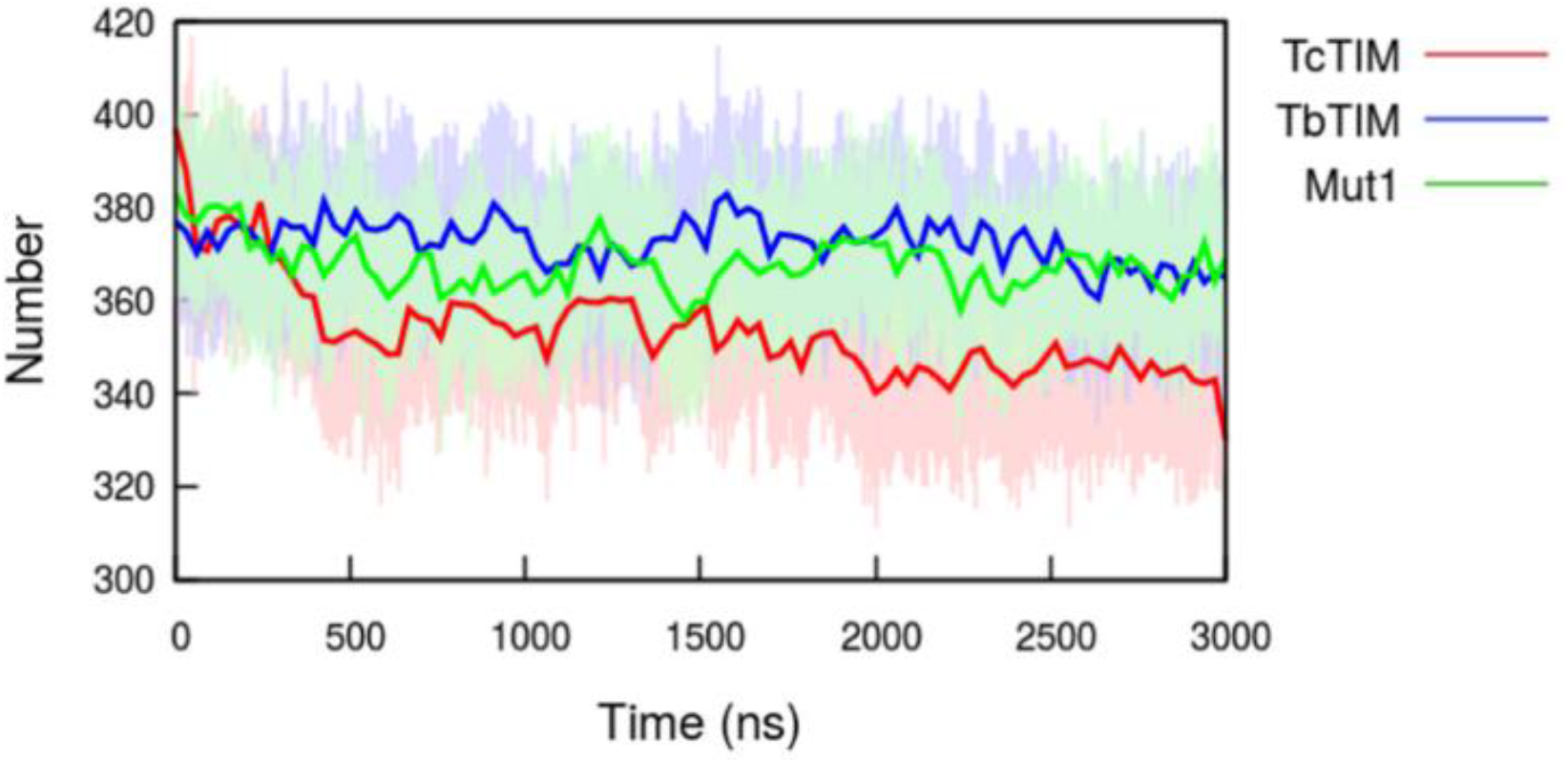
Total number of intramolecular hydrogen bonds in each simulation. Solid lines are Bézier curves that interpolate the data. Hydrogen bonds were defined with a cutoff radius of 0.3 nm for atoms whose acceptor-hydrogen-donor angle is greater than 120°^59^.

Species-specific inhibition of TIMs can be achieved by targeting a non-conserved amino acid (Cys15) that lies at the dimer interface and which is important for catalysis^11^. The susceptibility of TcTIM to thiol agents is approximately 40 times higher than that of TbTIM^60^. Figures S17 show the amino acids that form hydrogen bonds with Cys15 (Cys14 in TbTIM). Hydrogen bonds were determined with a cutoff angle of 30° for the hydrogen-donor-acceptor angle and a cutoff radius of 0.3 nm; OH and NH groups were regarded as donors, and O and N as acceptors. The residues that participate in hydrogen bonds with this cysteine change after the first microsecond in all three systems. In particular, those formed with residues Gly 73 and Ala 74 in TcTIM, with residue Ala 236 in TbTIM and with residues Phe 75 and Ser 80 in Mut1. Interestingly, TbTIM’s Cys14 interacts with residues in region 8 from the same monomer, while TcTIM and Mut1’s Cys15 interact with residues from region 3 in the other monomer. Fluctuations in the hydrogen bond network for Cys14/15 can only be noticed in simulations longer than 1 μs. Only the hydrogen bond between Cys15 monomer B and Phe75 monomer A in TcTIM was identified as such by RIP-MD^45^, the other interactions were identified as Cα contacts.

## DISCUSSION

### Biological relevance

TIM has four catalytic residues: Asn12, Lys 14, His 96 and Glu 168^2,10^. TcTIM is the only protein with cation-π interactions and one of them is between two of the catalytic residues: His 96 and Lys 14 (Fig. S2). Cation-π interactions can enhance binding energies by 2–5 kcal/mol, making them competitive with hydrogen bonds^61^. Another interaction only found in this protein occurs between catalytic Glu 168 and residue Arg 100, they form both a salt bridge and a hydrogen bond (Fig. S4, S5). In contrast, a salt bridge between catalytic Lys 14 (13 in TbTIM) and residue Glu 98 (97 in TbTIM) was observed in all three proteins (Fig. S4). This is a conserved salt bridge that has been observed in several crystal structures^29,30^. Lys 14 is also involved in the main network of hydrogen bonds in TcTIM. It forms a hydrogen bond with its neighbor, catalytic Asn 12, which in turn forms a hydrogen bond with residue Thr 76 from the other monomer and with the other catalytic residue, His 96 (Fig. S5). Similarly, Mut1 forms a hydrogen bond between two of the catalytic residues, Lys 14 and Asn 12, which in turn forms a hydrogen bond with residue 76 from the other monomer. However, unlike TcTIM, these residues are not connected to others in the network (Fig. S9). In both Mut1 and TbTIM, catalytic Asn 12 (Asn 11 for TbTIM) forms a hydrogen bond with residue Val 234 (Val 233 in TbTIM) and Lys 14 (Lys 13 in TbTIM) forms one with Gly 236 (Gly 235 in TbTIM) (Fig. S10, S11, S13, S15). Catalytic Glu 168 forms three hydrogen bonds in TcTIM: with Arg 100, Val 128 and Glu 130, but it only forms the last two bonds in TbTIM and Mut1 (Fig. S5, S11, S12, S14, S15).

Regions 1, 4 and 8 are known to become more rigid when the dimer is formed^30^. Region 1 forms several hydrogen bonds with region 3 of the opposite monomer. TcTIM forms six of these bonds at the interface, while TbTIM and Mut1 form only two (Fig. S5, S6, S9). Of all regions, region 4 had the highest number of salt bridges, comprising 13 residues in TcTIM, 11 in TbTIM and 10 in Mut1 (Fig. S4). Salt bridges between Arg 99 (Arg 98 in TbTIM) and Glu 78 (Glu 77 in TbTIM) involving both monomers were found at the interface of all three proteins. Only TcTIM formed salt bridges in region 8, two in each monomer.

The same salt bridge between residues Arg 192 (Arg 191 in TbTIM) and Asp 228 (Asp 227 in TbTIM) was observed in the two native proteins but not in Mut1 (Fig. S4). Since it has been shown that this conserved bridge is important for the efficient folding of TIM^62^, we expect that Mut1 will have a low recovery of activity in denaturation and refolding experiments.

TcTIM had a high RMSF value in loops 5 and 6 monomer A, but a low value in the same loops in monomer B (Fig. 3). Five amino acids in monomer B are involved in salt bridges, but only one in monomer A. Three amino acids from region 6 form salt bridges in monomer A, and four in monomer B. One of them is catalytic Glu 168. Similarly, TbTIM forms more salt bridges in monomer A loop 5, than in monomer B. Six residues from loop 5 monomer A form salt bridges but only three in monomer B. Only one residue in loop 6 monomer B forms a salt bridge (Fig. S4). Mut1 has a similar number of residues from loop 5 involved in salt bridges in both monomers, which explains why neither one of them has a high RMSF value. Salt bridges can vary in strength from weak (0.5 kcal/mol) to strong (3–5 kcal/mol) and play an important role in structure stabilization^63^. This may explain why the chimeric protein has a lower catalytic efficiency than its parent protein^14^.

The RMSF of TcTIM residues (Fig. 3) showed low values for the end of each monomer. This is explained by the network of hydrogen bonds (Fig. S7, S8) Amino acids at the end of the chain are connected to other residues forming a chain of hydrogen bonds that is not observed in the other two proteins. Furthermore, TcTIM has more interactions between monomers than the other two proteins. Some of these interactions were found in the mutant but not on TbTIM. This may explain why TcTIM has higher thermal stability even though it forms less hydrogen bonds than TbTIM^26^.

Based on this, we expect that Mut1 would have a thermal stability higher than TbTIM but lower than TcTIM.

### The need for long simulations

While the RMSD helped to identify the need to increase the simulation time of all three systems, it is not the only quantity where this issue can be noticed. When the RMSF of the first 200 ns of the simulation is compared with the RMSF of the last 200 ns of each simulation, noticeable differences can be observed (Fig. 9, S18, S19). One of the two main peaks in TcTIM and in the RMSF of TbTIM do not even appear at the beginning of the simulations. Furthermore, fluctuations in many amino acids decrease, which help to highlight the relevance of the main peaks. In contrast, when one compares the last 200 ns of the simulation with the last microsecond of the trajectory, differences are considerably smaller. An exception to this appears in loop 6 monomer B of TbTIM, which is only observed when the average is calculated over the last microsecond of the simulation.

**Figure 9.**
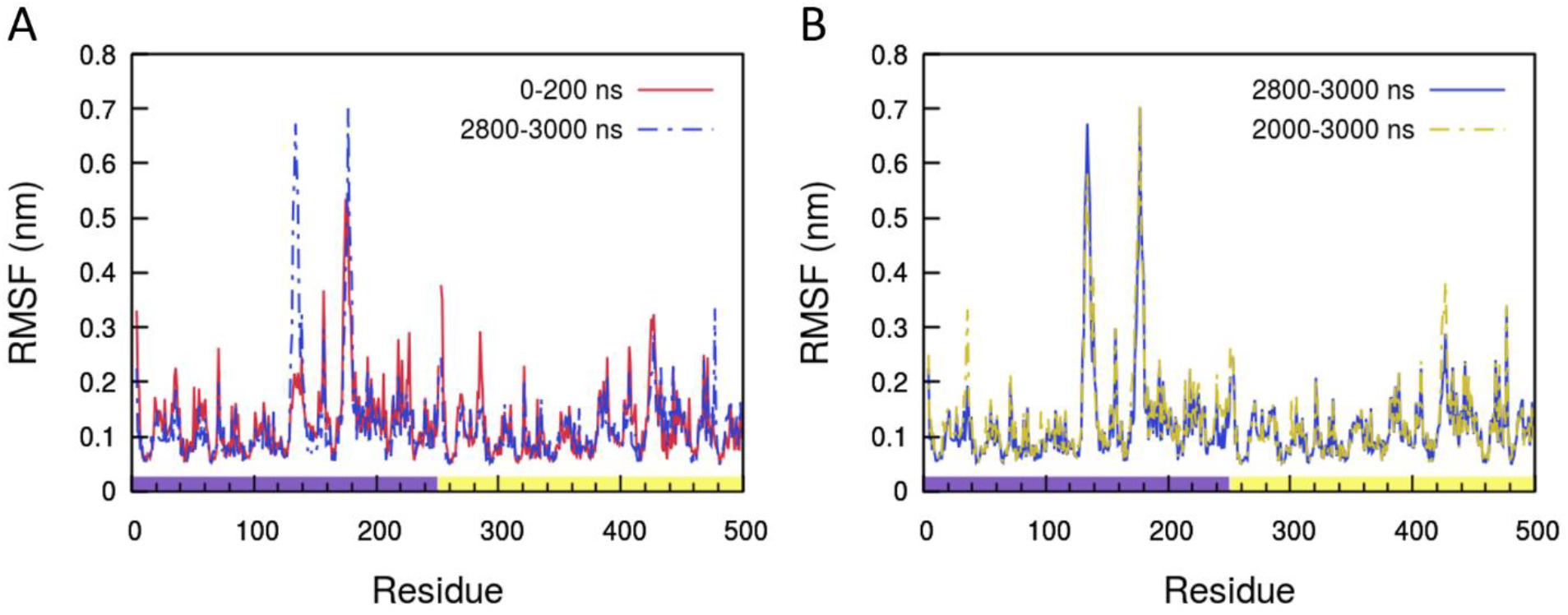
Changes over time in the root mean square fluctuations of the residues of TcTIM. A) RMSF of the first 200 ns of the simulation vs the last 200 ns, and B) RMSF of the last microsecond of the trajectory vs the last 200 ns. The peak in loop 5 of monomer A does not appear at the beginning of the trajectory and fluctuations in the minor peaks decrease at longer times. The color bar at the bottom of figure B distinguishes the residues in monomer A (purple) from those in monomer B (yellow).

One way to monitor convergence of simulations is to plot the average of a quantity over different time intervals. If the average changes with different time windows, then the system is not properly equilibrated. Figure 10 shows the number of intramolecular hydrogen bonds for the TcTIM trajectory. Horizontal lines mark the averages taken over different time windows. The average for the first 500 ns is 7% higher than that of the last 500 ns. The average over the last microsecond of the trajectory equals the average over the last 500 ns (345 hydrogen bonds), which indicates that the trajectory has stabilized over the last microsecond.

**Figure 10.**
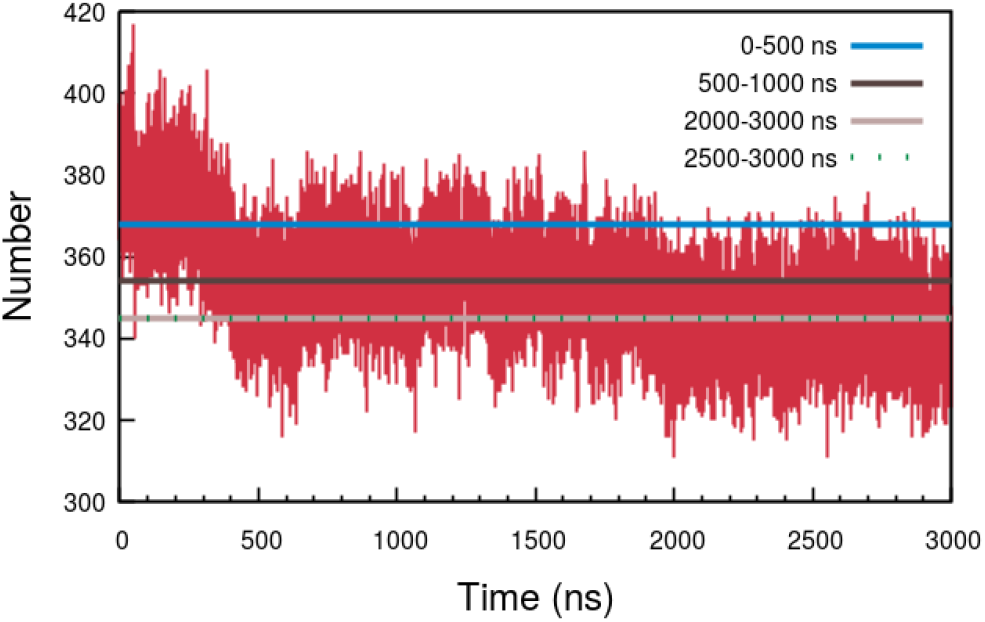
Number of hydrogen bonds in TcTIM. Convergence (horizontal lines) is reached around 2 μs. Averaging over the last 500 ns or over the last microsecond of the simulation produces the same average (345 hydrogen bonds; the lines overlap).

In order to find how the hydrogen bond networks changed with time, we used the Gromacs built-in cluster tool to generate clusters for the first and last 500 ns of the simulations. The clusters were generated using the gromos method with a RMSD cutoff of 0.2 nm. We then used RIP-MD^46^ to analyze the RINs in the representative structure of the most populated cluster. Figures S20 and S21 show the most significant changes in TcTIM. At the beginning of the simulation there is little connectivity between the main hydrogen bond networks of each monomer, when residues 75, 78 and 99 in monomer B interact with residues 15 and 103 in monomer A, these two networks merge into one (Fig. 11).

**Figure 11.**
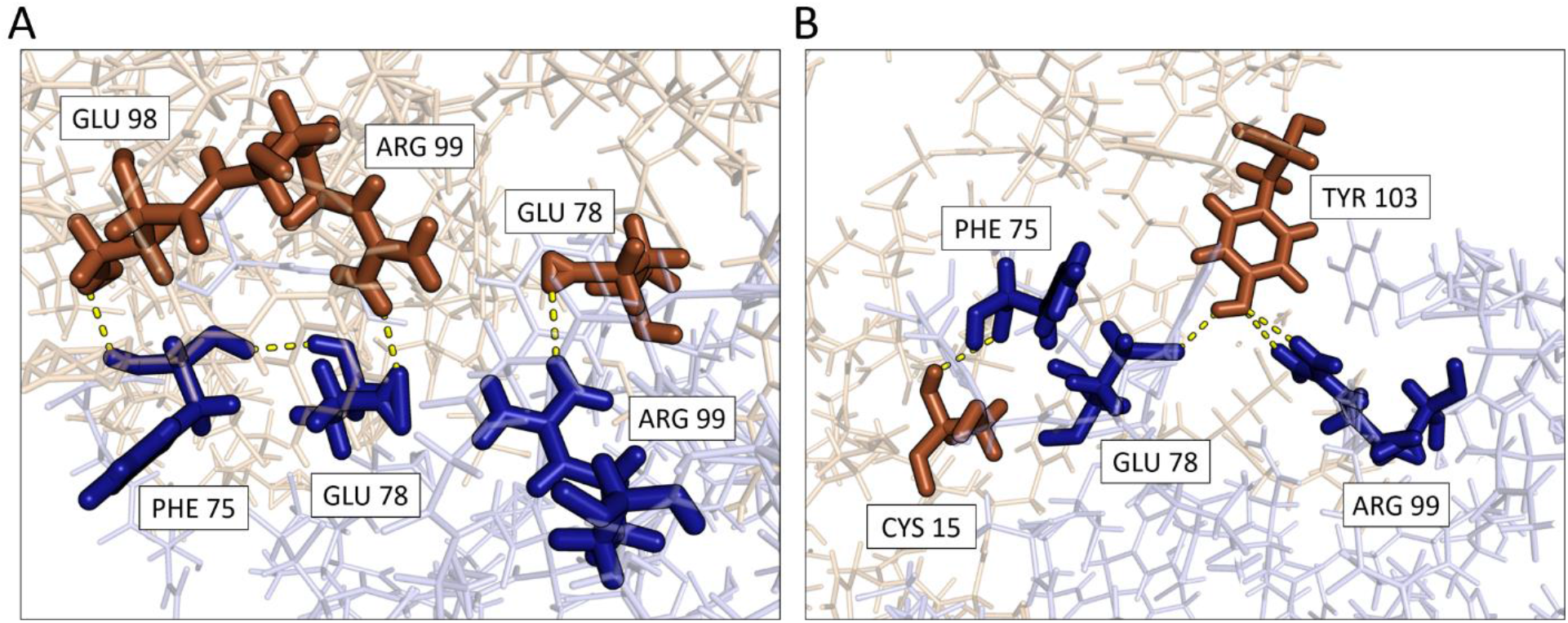
Changes in the hydrogen bond network of TcTIM. Interface between monomers (A) at the beginning and (B) at the end of the simulation. Residues Phe 75, Glu 78 and Arg 99 in monomer B (blue) change their interaction with residues in monomer A (orange).

Changes in protein dynamics are in turn reflected in the interactions between amino acids. Some interactions appeared after the first 500 ns of the simulation, *e.g.,* the π-π interaction between residues 187 and 210 in monomer A of TbTIM, and others became more stable only after the first microsecond of the simulation, *e.g.,* the π-π interaction between residues 36 and 224 of monomer B in TcTIM (Fig. 12).

**Figure 12.**
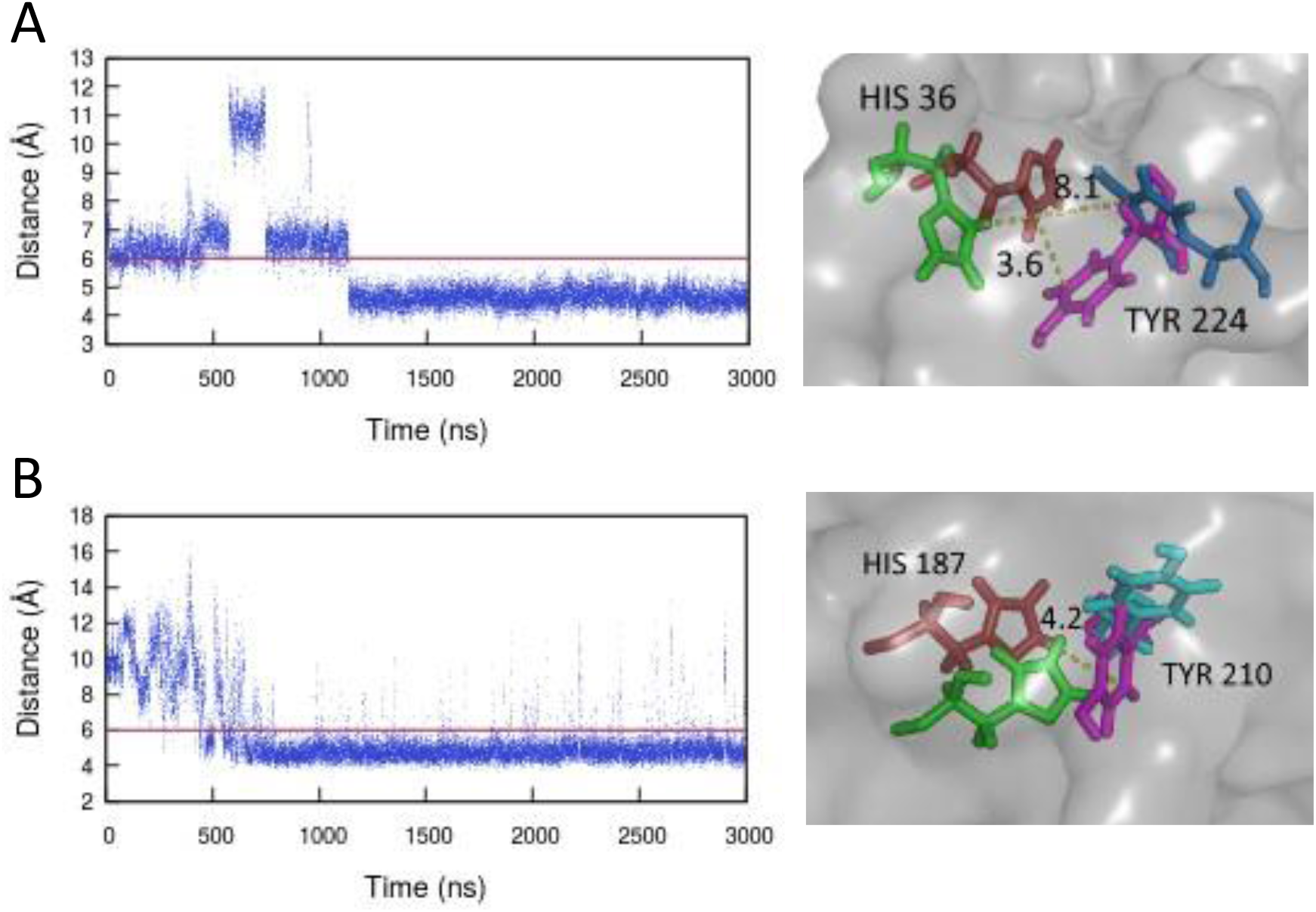
π-π interactions in TcTIM between residues His 36 and Tyr 224 of monomer B (A) and in TbTIM between residues His 187 and Tyr 210 in monomer A (B). A red line marks the threshold that defines the π-π interaction. At the right side of each graph, conformation changes between the first frame (green and blue) and the last frame (red and purple) of each simulation are shown. The interaction in TcTIM was not observed during the first 500 ns of the simulation. The interaction in TbTIM was observed since the beginning but it only become stable after the first microsecond.

## CONCLUSIONS

We have performed molecular dynamics simulations on three different TIM proteins: TcTIM, TbTIM and a chimeric protein, Mut1. We examined the different electrostatic interactions that occur in these proteins: salt bridges, cation-π interactions, π-π interactions and hydrogen bonds, and also explored the impact of simulation length on them. Some of these interactions appeared only after the first microsecond of the simulation, and convergence of the number of hydrogen bonds was only reached in the last of the 3 μs of the simulation. Although TcTIM forms less hydrogen bonds than TbTIM and Mut1, they form a network that spans almost the entire protein, connecting the residues in both monomers. Key differences were found in the interactions that the catalytic amino acids form, such as a cation-π interaction between catalytic amino acids Lys 14 and His 96, only observed in TcTIM, but a salt bridge between catalytic residue Lys 14 and Glu 98 was observed in all three proteins. Further experiments will be required to confirm our hypothesis on the thermal stability of Mut1.

## Supporting Information

Figure S1: Minimum distance between loops 6 and 7, Figure S2: Cation-π interactions for TcTIM, Figure S3: π-π interactions for TcTIM, TbTIM and Mut, Figure S4: Salt bridges for TcTIM, TbTIM and Mut, Figures S5-S16: Hydrogen bond networks in the different systems, Figure S17: Hydrogen bonds in Cys 14/15 monomer, Figures S18-S19: Changes over time in the root mean square fluctuations (RMSF), Figure S20: Main hydrogen bond networks for TcTIM in the most populated cluster of the first 500 ns of the simulation. Figure S21: Hydrogen bond networks for TcTIM in the most populated cluster of the last 500 ns of the simulation.

## AUTHOR INFORMATION

### Author Contributions

The manuscript was written through contributions of all authors. All authors have given approval to the final version of the manuscript.

## ACKNOWLEDGMENT

The authors thank Dr. Mónica Rodríguez-Bolaños and Dr. Ruy Perez-Montfort for fruitful discussions. CCG thanks the Province of Ontario Trillium Scholarship Program. MK thanks the Natural Sciences and Engineering Research Council of Canada (NSERC) and the Canada Research Chairs Program. Computing facilities were provided by SHARCNET (www.sharcnet.ca), Compute Canada (www.computecanada.ca).

## ABBREVIATIONS

TIM: triosephosphate isomerase
TcTIM: *Trypanosoma cruzi*
TbTIM: *Trypanosoma brucei*
MD: molecular dynamics
PDB: Protein Data Bank
RSMF: root-mean-square fluctuation
PME: particle-mesh Ewald method
P-LINCS: Parallel Linear Constraint Solver
RMSD: root-mean-square deviation.

