## Supplementary Information for "Highly similar sequence and structure yet different biophysical behaviour: A computational study of two triosephosphate isomerases"

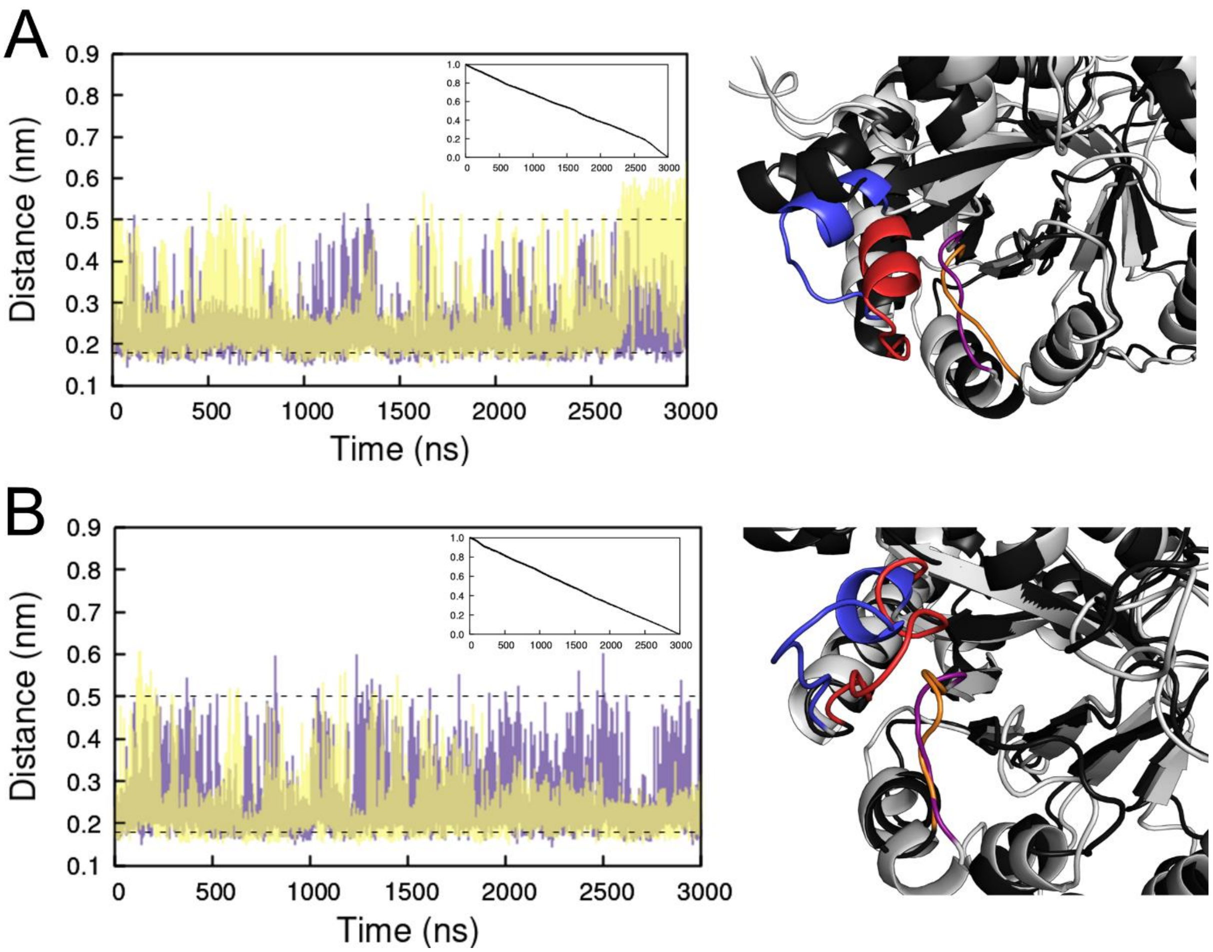


Fig. S1. Minimum distance between loops 6 and 7 in TbTIM (A) and Mut1 (B) for monomer A (purple) and monomer B (yellow). Dashed lines mark the two different states sampled by the loops. Inside: cross-correlation between the loop state of the two monomers. Right: alignment of the open (gray) and closed (black) conformations. Loop 6 is shown in red for the closed conformation and blue for the open conformation, loop 7 is shown in orange (closed) and purple (open).

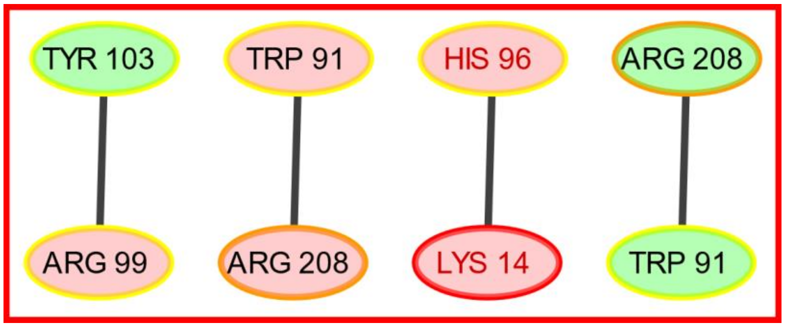


Fig. S2. Cation-π interactions for TcTIM throughout the last 2 μs of the simulation. Amino acids in monomer A are shown in green and residues in monomer B in red. Each node is coloured according to the color scheme for regions in Fig. 1 of the main text. The catalytic residues are written with red text.

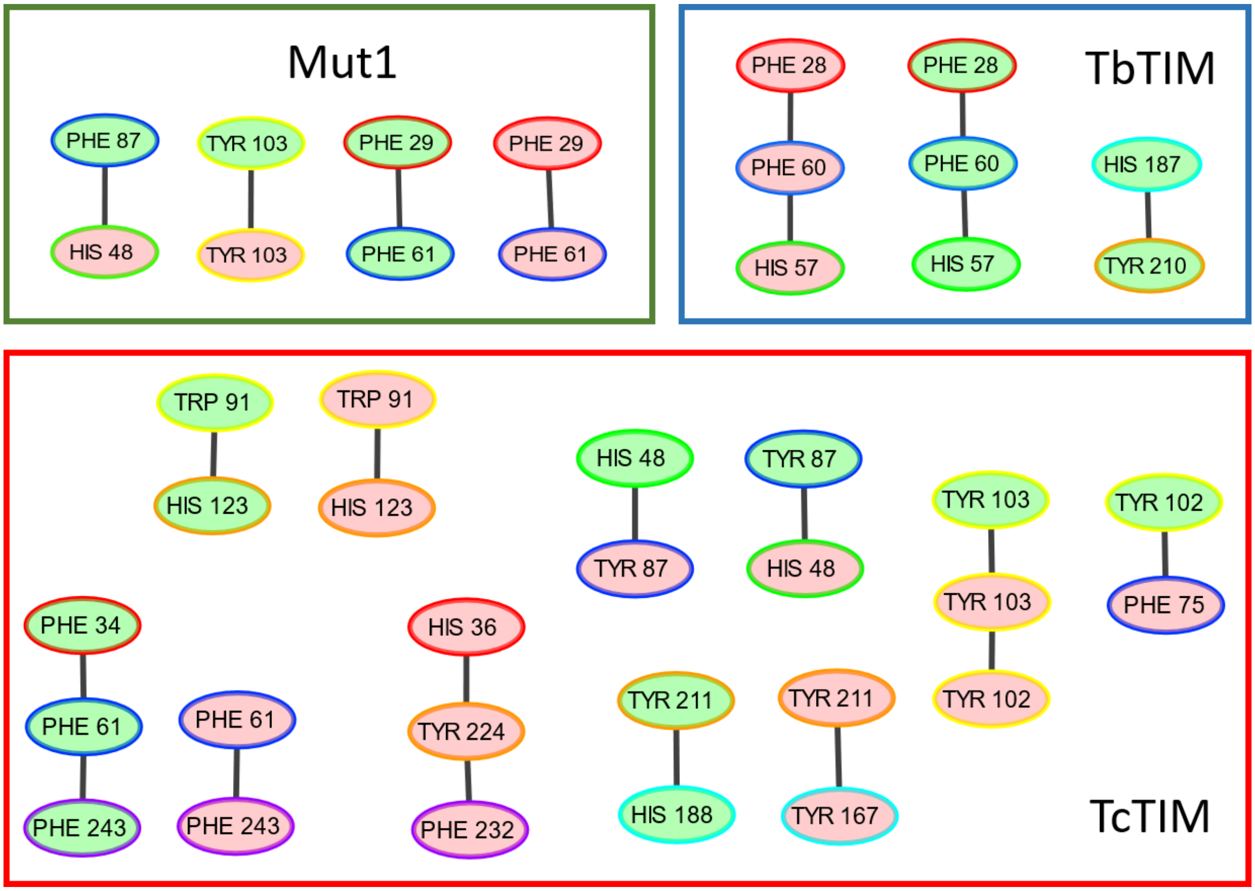


Fig. S3. π-π interactions for TcTIM, TbTIM and Mut1 throughout the last 2 μs of the simulations. Amino acids in monomer A are shown in green and residues in monomer B in red. Each node is coloured according to the color scheme for regions in Fig. 1 of the main text.

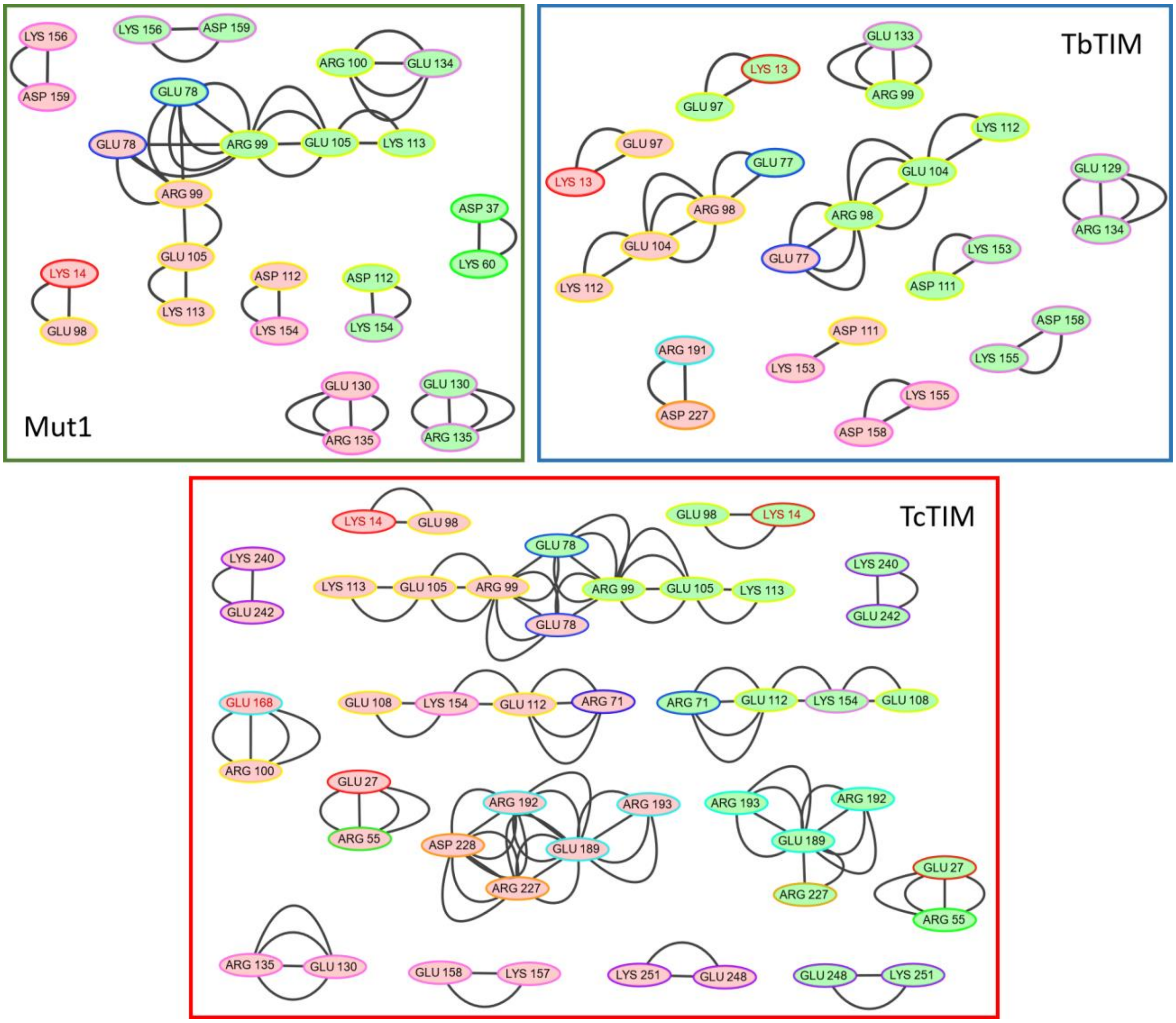


Fig. S4. Salt bridges for TcTIM, TbTIM and Mut1 throughout the last 2 μs of the simulations. TcTIM forms more salt bridges than the other two proteins. Amino acids in monomer A are shown in green and residues in monomer B in red. Each node is coloured according to the color scheme for regions in Fig. 1 of the main text. The catalytic residues are written with red text.

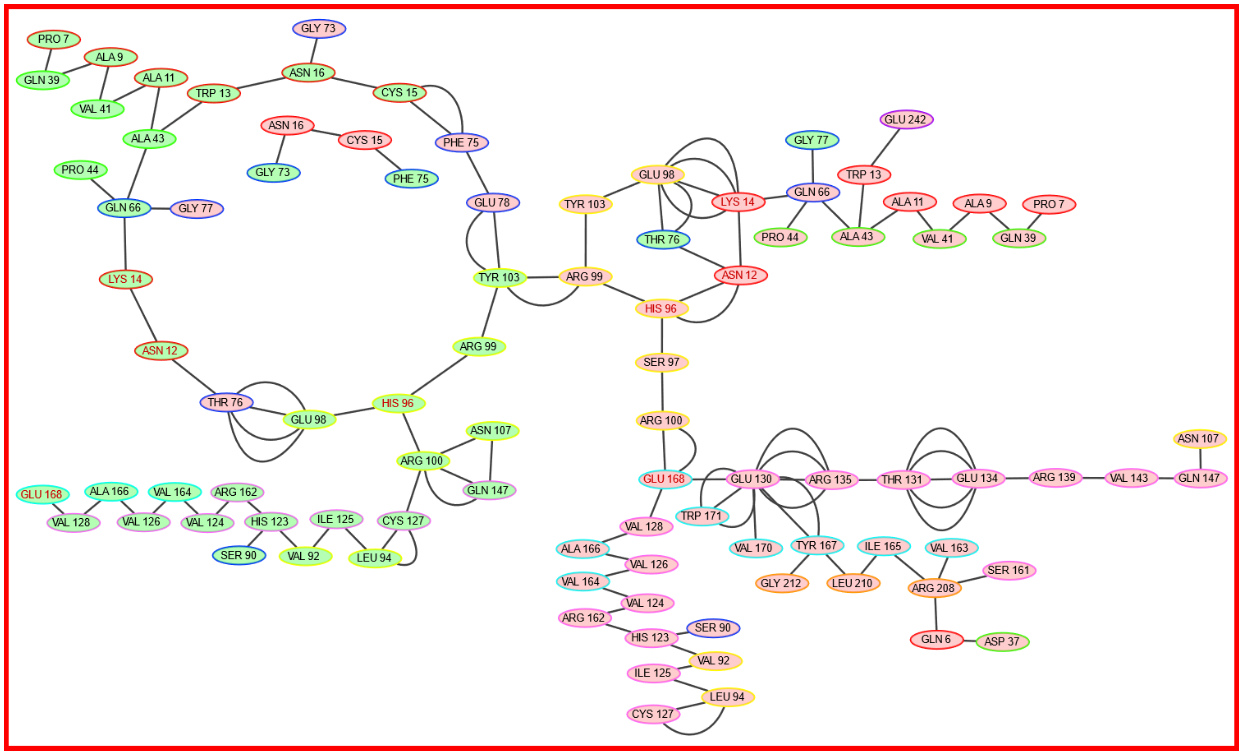


Fig. S5. Main hydrogen bond network in TcTIM throughout the last 2 μs of the simulation. Each node is coloured according to the color scheme for regions in Fig. 1 of the main text. The catalytic residues are written with red text.

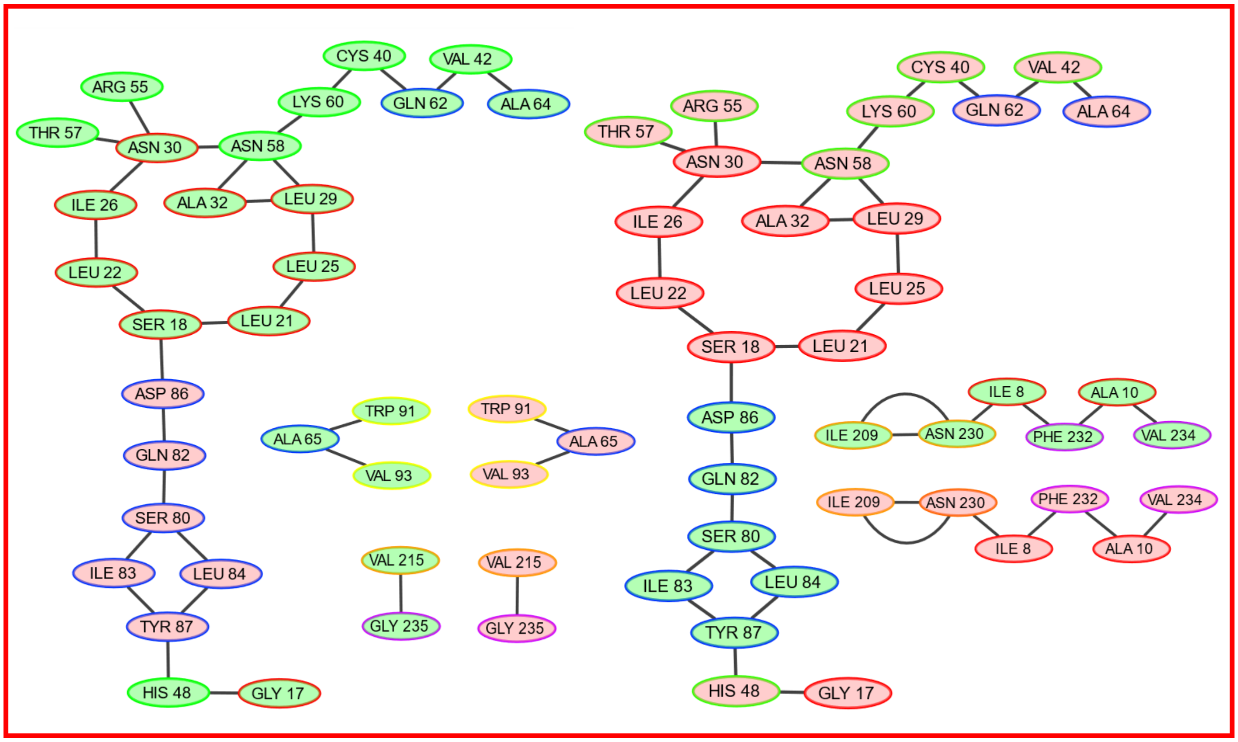


Fig. S6. Hydrogen bonds in TcTIM throughout the last 2 μs of the simulation. These hydrogen bonds are found in both monomers. Amino acids in monomer A are shown in green and residues in monomer B in red. Each node is coloured according to the color scheme for regions in Fig. 1 of the main text.

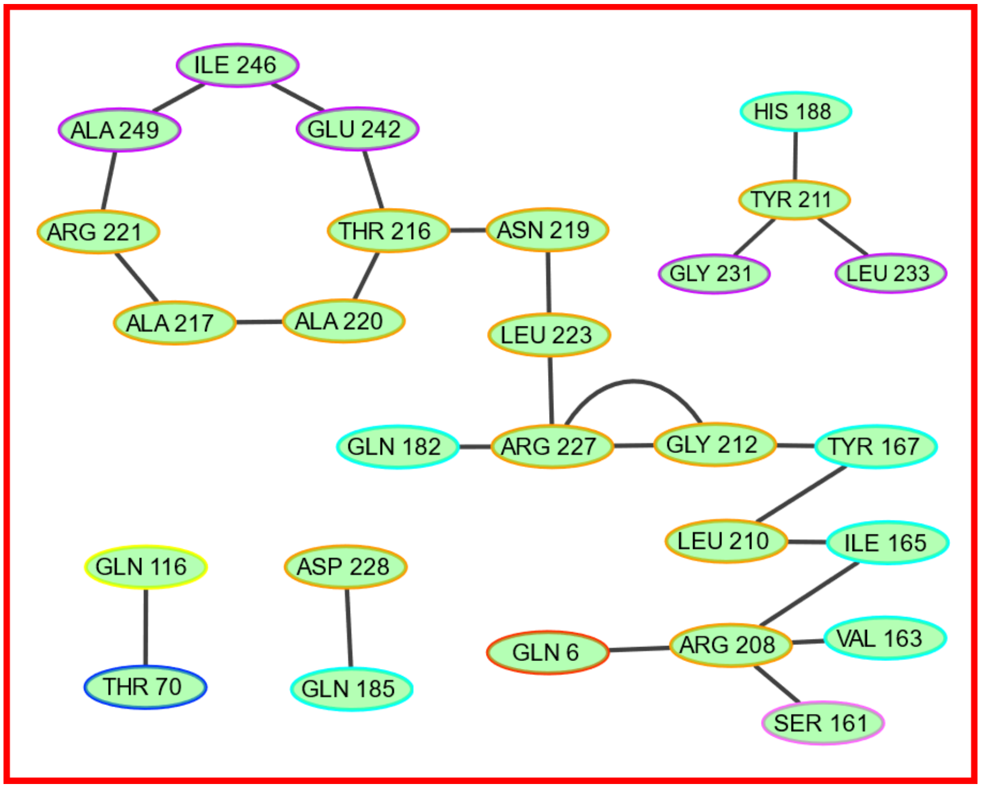


Fig. S7. Hydrogen bonds in monomer A throughout the last 2 μs of the TcTIM simulation. Each node is coloured according to the color scheme for regions in Fig. 1 of the main text.

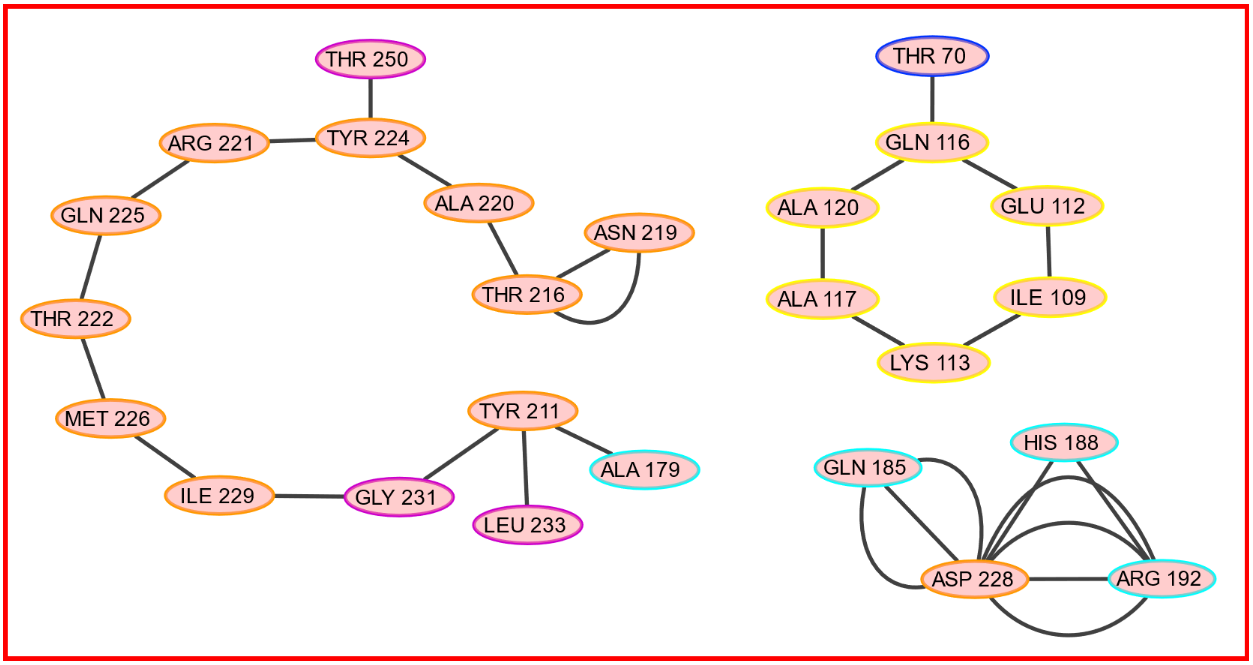


Fig. S8. Hydrogen bonds in monomer B throughout the last 2 μs of the TcTIM simulation. Each node is coloured according to the color scheme for regions in Fig. 1 of the main text.

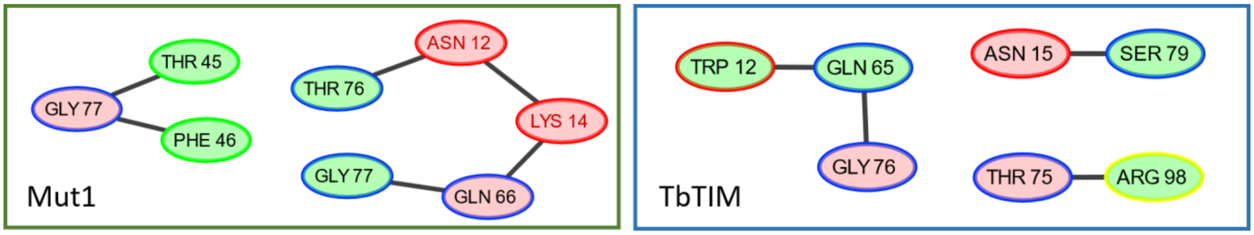


Fig. S9. Hydrogen bonds involving amino acids at the interface between monomers in Mut1 and TbTIM throughout the last 2 μs of the simulations. Amino acids in monomer A are shown in green and residues in monomer B in red. Each node is coloured according to the color scheme for regions in Fig. 1 of the main text. The catalytic residues are written with red text.

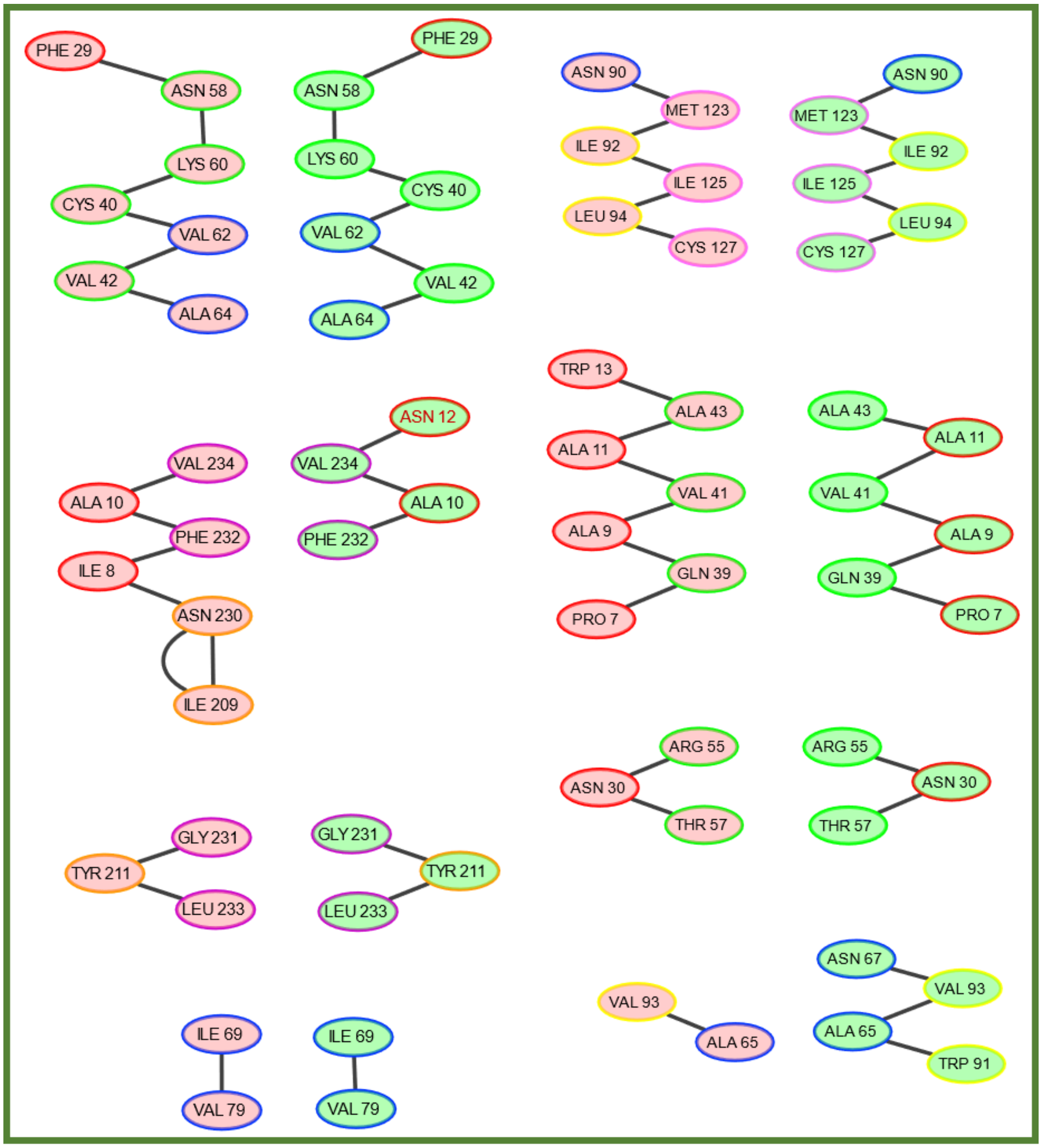


Fig. S10. Hydrogen bonds in Mut1 throughout the last 2 μs of the simulation. These hydrogen bonds are found in both monomers. Amino acids in monomer A are shown in green and residues in monomer B in red. Each node is coloured according to the color scheme for regions in Fig. 1 of the main text. The catalytic residue is written with red text.

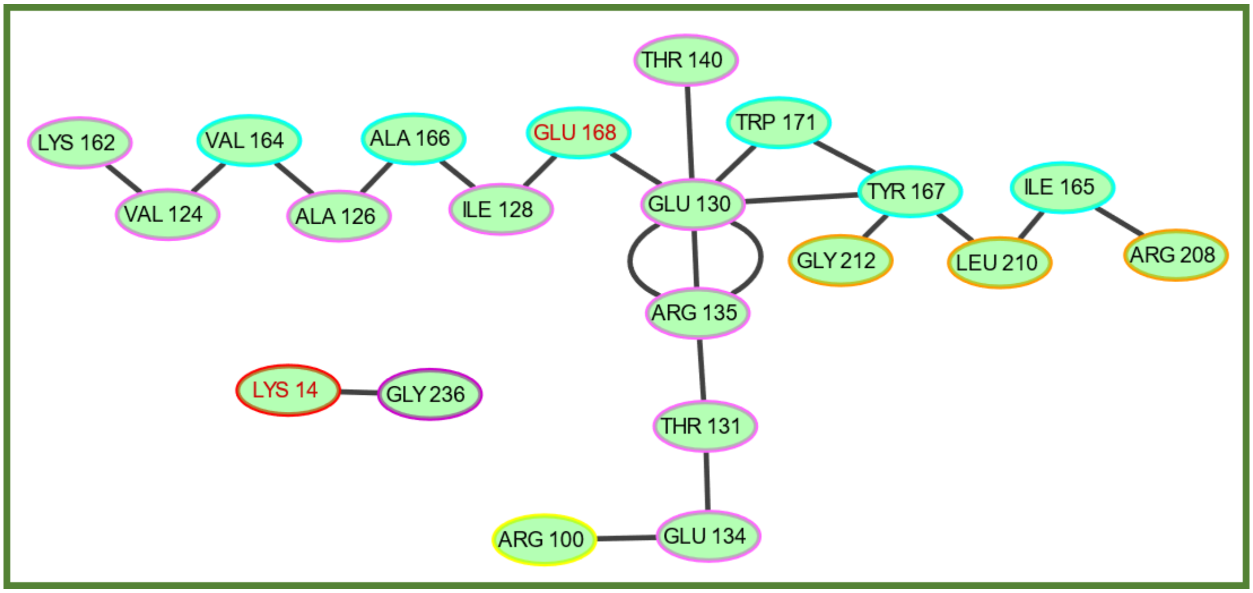


Fig. S11. Hydrogen bonds in monomer A throughout the last 2 μs of the Mut1 simulation. Each node is coloured according to the color scheme for regions in Fig. 1 of the main text. The catalytic residues are written with red text.

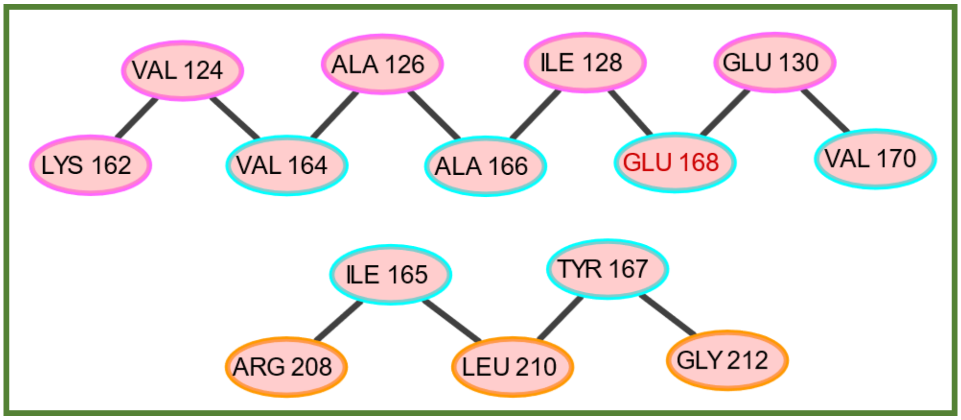


Fig. S12. Hydrogen bonds in monomer B throughout the last 2 μs of the Mut1 simulation. Each node is coloured according to the color scheme for regions in Fig. 1 of the main text. The catalytic residue is written with red text.

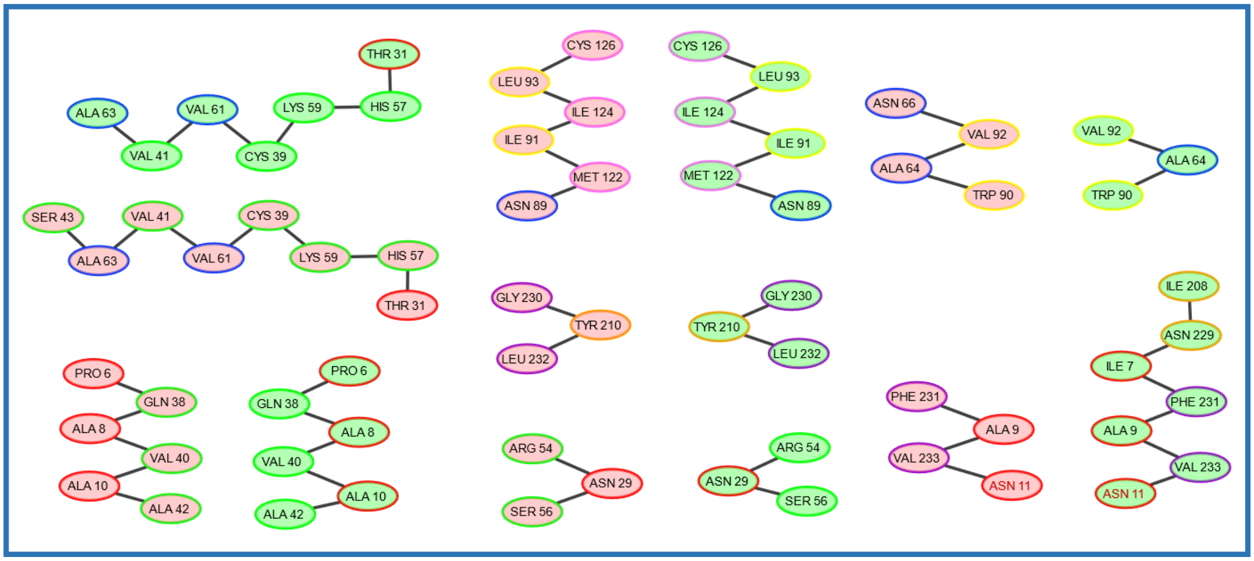


Fig. S13. Hydrogen bonds in TbTIM throughout the last 2 µs of the simulation. These hydrogen bonds are found in both monomers. Amino acids in monomer A are shown in green and residues in monomer B in red. Each node is coloured according to the color scheme for regions in Fig. 1 of the main text. The catalytic residues are written with red text.

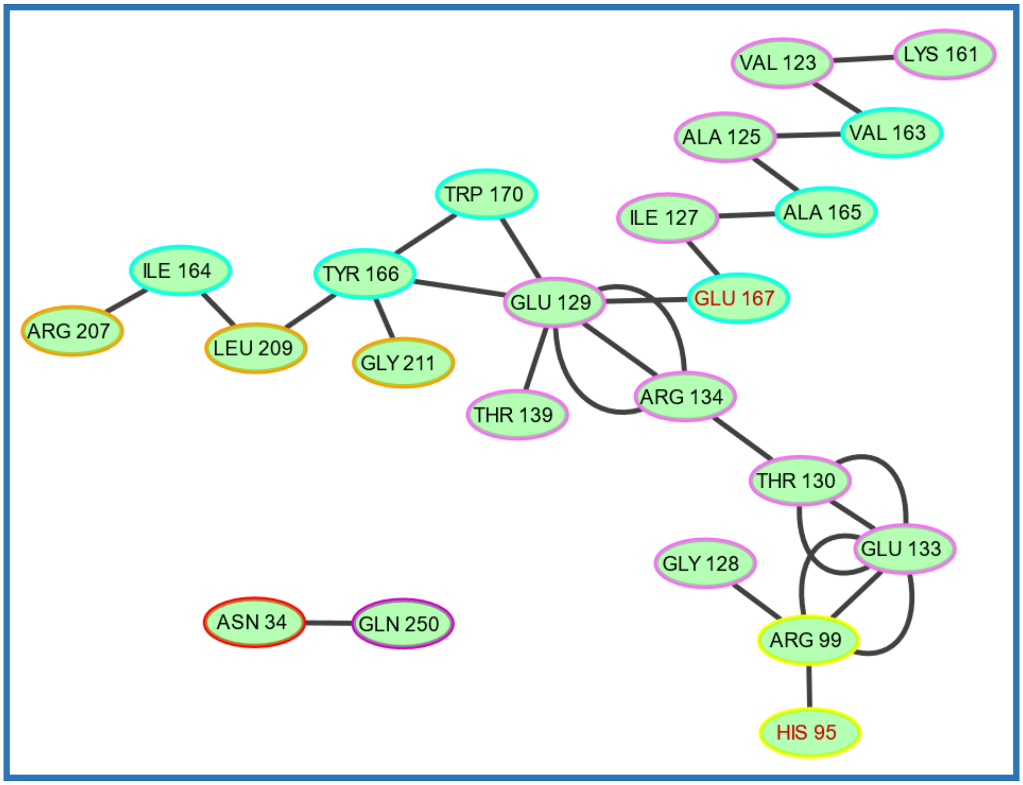

Fig. S14. Hydrogen bonds in monomer A throughout the last 2 µs of the TbTIM simulation. Each node is coloured according to the color scheme for regions in Fig. 1 of the main text. The catalytic residue is written with red text.

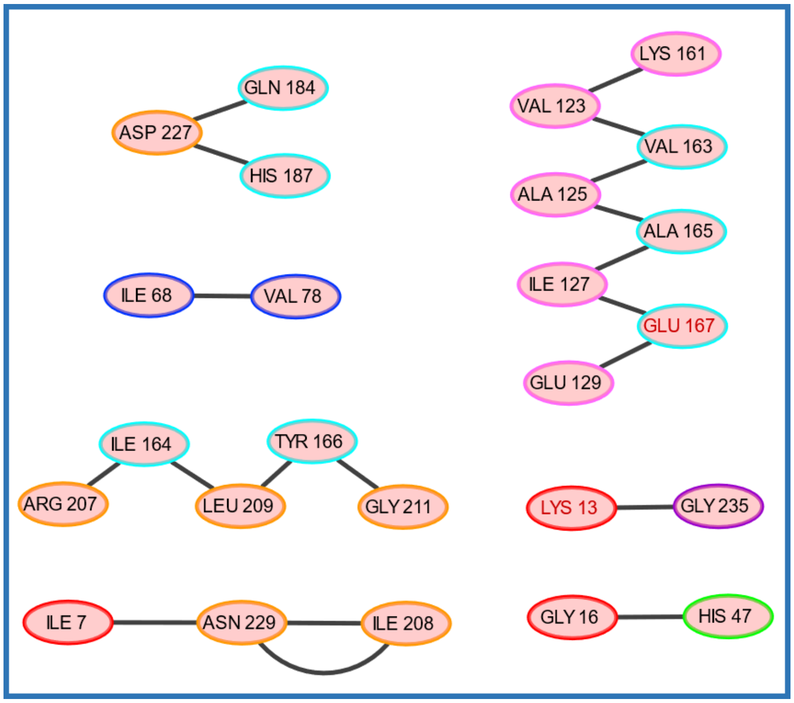


Fig. S15. Hydrogen bonds in monomer B throughout the last 2 μs of the TbTIM simulation. Each node is coloured according to the color scheme for regions in Fig. 1 of the main text. The catalytic residues are written with red text.

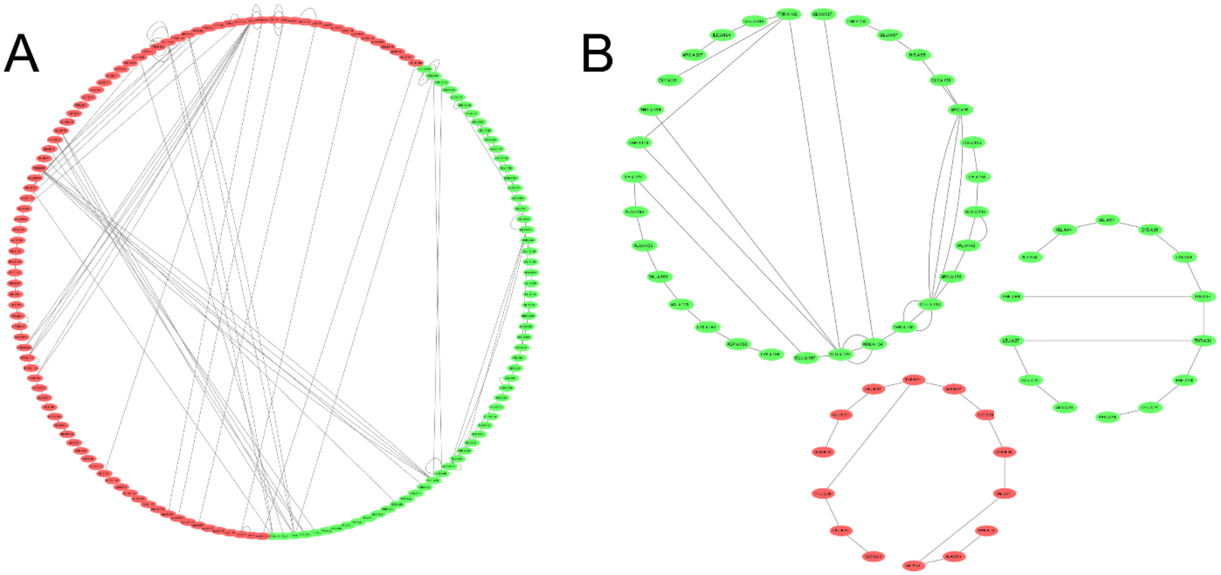


Fig. S16. Main hydrogen bond networks throughout the last 2 μs for A) TcTIM and B) TbTIM. Amino acids in monomer A are shown in green and residues in monomer B in red. Hydrogen bonds in TcTIM connect amino acids in a network that involves many interactions between monomers and extends throughout the whole protein, in contrast with TbTIM, whose networks are contained within each monomer and involve fewer residues.

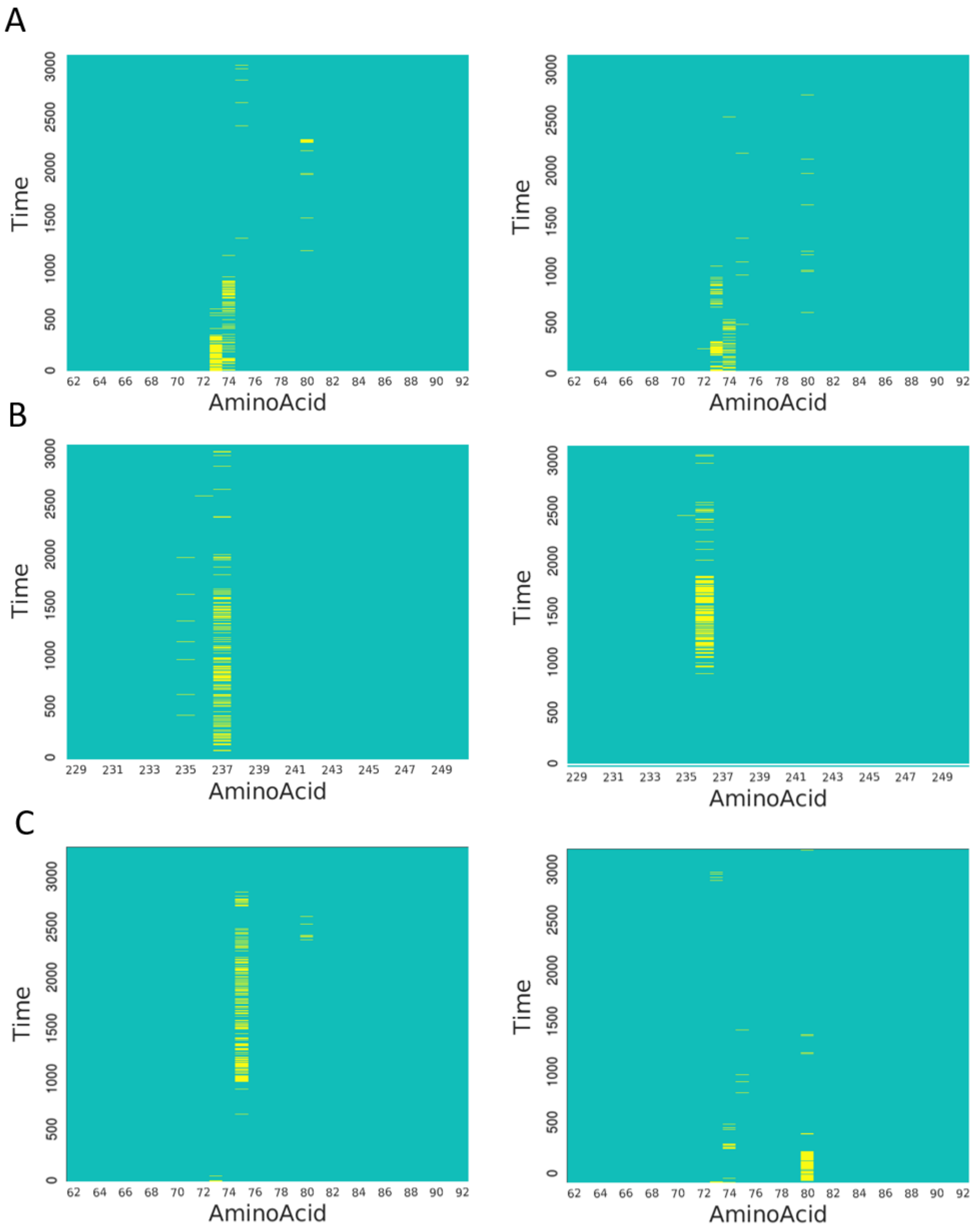


Fig. S17. Hydrogen bonds in Cys 14/15 monomer A (left) and monomer B (right) in: A) TcTIM, B) TbTIM and C) Mut1. This cysteine forms hydrogen bonds with region 3 of the other monomer in TcTIM and Mut1 and forms hydrogen bonds in region 8 of the same monomer in TbTIM.

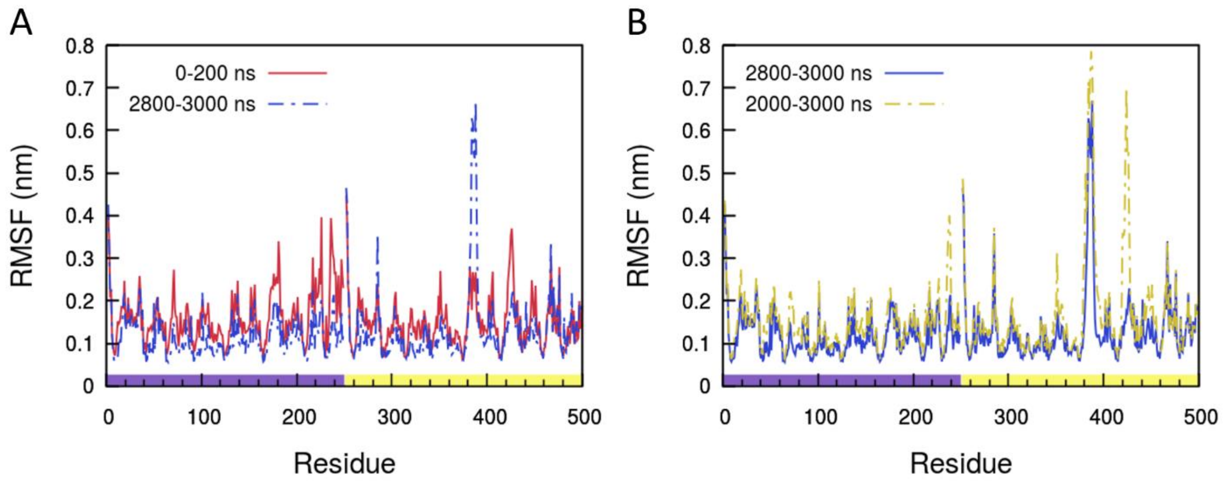


Fig. S18. Changes over time in the root mean square fluctuation (RMSF) of the TbTIM simulation. RMSF of the first 200 ns of the simulation vs the last 200 ns (A), and RMSF of the last microsecond of the trajectory vs the last 200 ns (B). The color bar at the bottom of figure B distinguishes the residues in monomer A (purple) from those in monomer B (yellow).  There are no significant peaks in the RMSF of the beginning of the trajectory. The last 200 ns of the simulation failed to capture the peak at loop 6 monomer B.

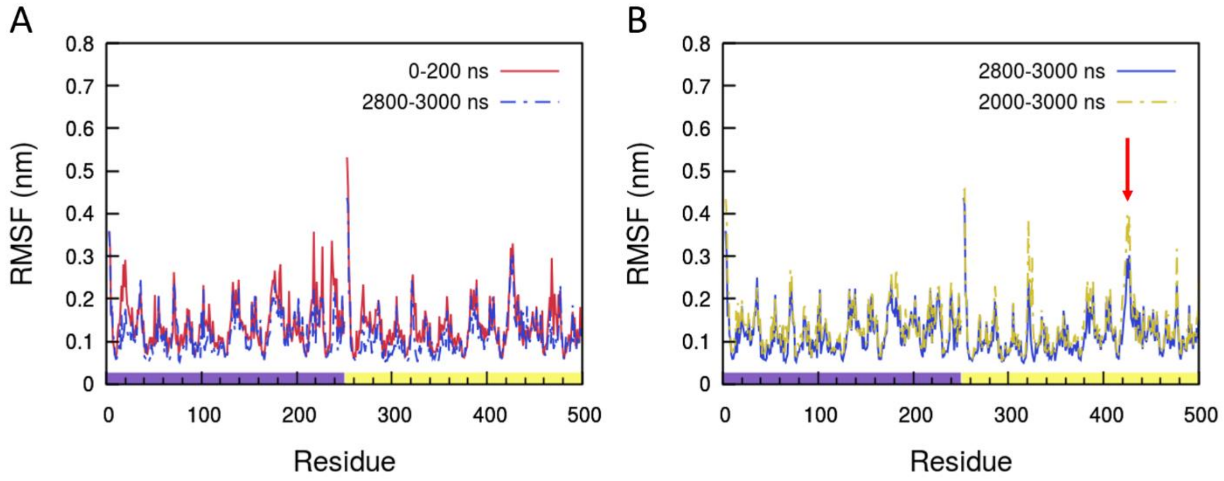


Fig. S19. Changes over time in the root mean square fluctuation (RMSF) of the Mut1 simulation. RMSF of the first 200 ns of the simulation vs the last 200 ns (A), and RMSF of the last microsecond of the trajectory vs the last 200 ns (B). The color bar at the bottom of figure B distinguishes the residues in monomer A (purple) from those in monomer B (yellow). Fluctuations at the minor peaks decreased with time and the peak at loop 6 monomer B (red arrow) increased.

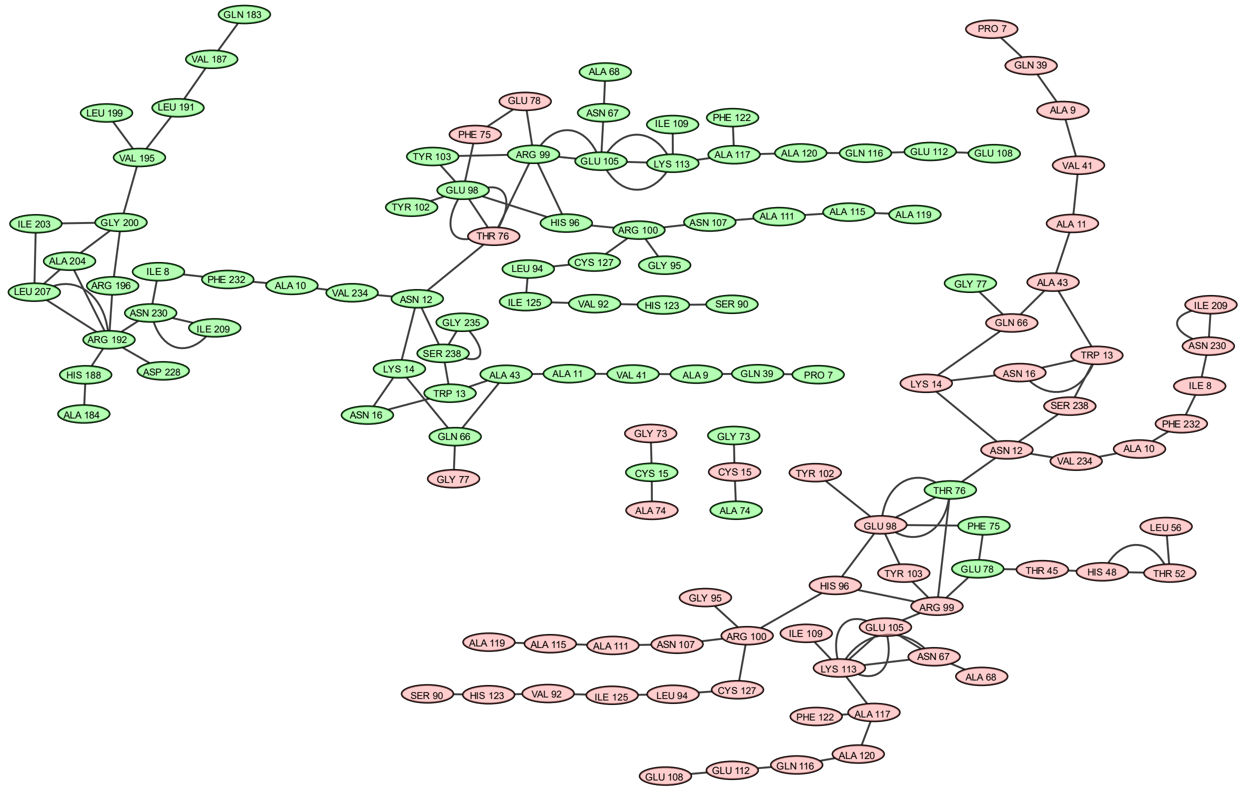


Fig. S20. Main hydrogen bond networks for TcTIM in the most populated cluster of the first 500 ns of the simulation. There is a different network for each monomer. Amino acids in monomer A are shown in green and residues in monomer B in red.

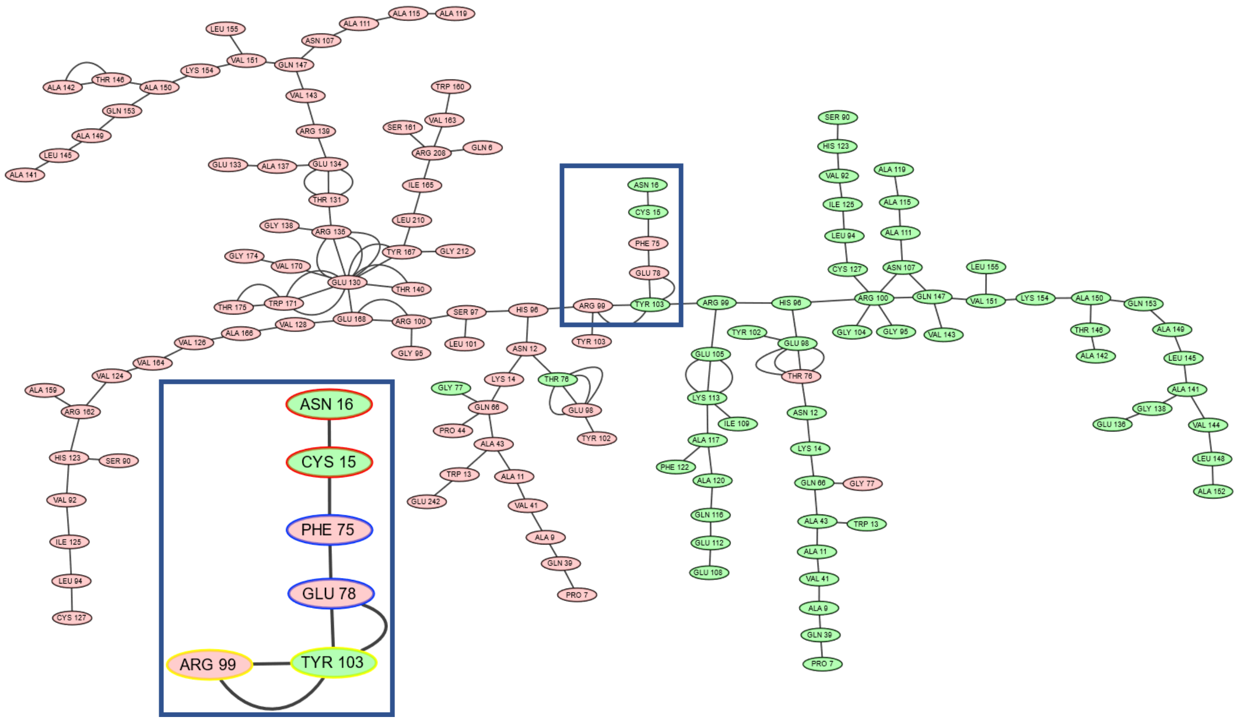


Fig. S21. Hydrogen bond networks for TcTIM in the most populated cluster of the last 500 ns of the simulation. The residues in both monomers are connected through a single network of hydrogen bonds. Highlighted in blue are the residues at the interface of the hydrogen bond network (Fig. 11 in the main text). Amino acids in monomer A are shown in green and residues in monomer B in red.
